# Global genomic epidemiology of *Candida auris*: analysis of 12,644 whole genome sequences from 1997-2024

**DOI:** 10.64898/2026.02.03.703534

**Authors:** Hugh Gifford, Nicolas Helmstetter, Hanane Zerrouki, Duncan Wilson, Johanna Rhodes, Rhys A. Farrer

## Abstract

*Candida auris* is a critical priority fungal pathogen (World Health Organization) that has emerged into human populations pre-1996 from an unknown environmental reservoir. Genomic sequencing data has been used extensively in the last decade, leading to a largely unharnessed dataset with potential to unlock understanding of emerging fungal pathogen evolution. Here, we compiled publicly available Illumina paired-end whole genome sequences (WGS) for variant calling, supplemented by isolates from the world’s first four-clade single-facility outbreak in Algeria (*n* = 7), totaling 12,644 WGS. We describe the geographic, clinical, and temporal epidemiology of the outbreak across the globe between 1997 and 2024, indicating a conserved six-clade structure. Despite the evidence of earlier species diversity in low- and lower-middle income countries (LLMIC), the majority of WGS derived from high-income (12,035, 93.2%) and upper-middle-income countries (719, 5.57%), where cases are believed to be imported from endemic regions, with few from lower-middle-income countries (165, 1.28%) and none from lower-income countries. Copy number variation was present, including azole drug target *ERG11*, with seven isolates displaying ten to fifteen copies. Alarmingly, the standing variation of *C. auris* reveals emerging variation in hot-spots of the echinocandin drug target *FKS1*, which encodes a β-1,3-glucan synthase. Eighteen emerging *FKS1* hot-spot variants have not been detected or described in databases, and mostly occured in Northern America (57/61, 93.4%), where echinocandin monotherapy is standard treatment. Well-studied *FKS1* variants known to cause resistance are significantly more common in clade I isolates derived from urine compared to blood, consistent with a role for the urinary niche as a low echinocandin concentration safe-haven for the development of resistance. The insights from this global genomic epidemiology survey of *C. auris* highlight sequencing inequality and detect ongoing genomic innovation in clinical settings, raising predictable and urgent concerns around the ongoing use of echinocandin monotherapy and potential emerging antifungal drug resistance-related genotypes in high-income settings.

## Introduction

*Candida auris*, also named *Candidozyma auris*^1,2^, is a global public health threat^3^ and critical priority human fungal pathogen^4^. *C. auris* is also classified as an urgent antimicrobial resistance (AMR) threat^5^, with species-wide fluconazole drug resistance exceeding >87%^6^. Since its first description in 2008^7^, hospital outbreaks have been reported across all WHO regions in more than 45 countries^8^, and are associated with prolonged hospitalisation (median 46-68 days) and high mortality (45%)^9^. Whole-genome sequencing (WGS) of more than 3,000 isolates has revealed six near-simultaneously emergent clades separated by at least 37,000 single nucleotide polymorphisms (SNPs)^10,11^. Despite the potential for molecular epidemiology to inform genetic drivers of resistance and virulence during the ongoing global outbreak^12,13^, analyses of the global genomic epidemiology of *C. auris* remain limited.

The first recorded four-clade single hospital outbreak to our knowledge took place in Tlemcen, Algeria^14^. Hypothetically, species genetic diversity may be associated with geographical proximity to ancestral habitats. The co-existence of three clades in a single country (clades I, III and IV in S. Africa) has previously been apportioned to introductions from other continents^15^. We have argued that assigning geographical origins to clades at present is likely to be misleading^16^. Indeed, the diversity of *C. auris* isolates in the African continent^17–19^ could represent proximity to ancestral strain habitats, but the paucity of sequencing in low- and lower-middle income countries (LLMIC) limits robust inference, obscures origin tracing, and drives global inequality^13^. Genomic sequencing, though largely generated in high-income settings, offers an opportunity to further resolve the global genomic epidemiology of *C. auris*. A critical priority in analysing the ongoing world outbreak is to survey genetic variation associated with drug resistance^20–22^.

Multiple genetic mechanisms drive drug resistance in *C. auris*. MARDy^23^, AFRbase^24^, FungAMR^25^ and CanDRes^26^ databases so far catalogue 130 variants across 29 *C. auris* genes that have been associated with resistance to the four main classes of antifungal drugs: azoles inhibiting lanosterol-14α-demethylase (fluconazole, voriconazole, itraconazole, isavuconazole, posaconazole, miconazole and ketoconazole), echinocandins inhibiting β-glucan synthase (anidulafungin, caspofungin, micafungin and rezafungin), polyenes binding cell membrane ergosterol (amphotericin B and nystatin), and antimetabolites (5-flucytosine). Resistance is most common against azoles (90%) and less common against polyenes (30%) and echinocandins (5%)^8^. Unlike for other pathogens of major global public health importance, such as *Mycobacterium tuberculosis* (TB), *Plasmodium falciparum* (Malaria) or HIV, the current policy for *C. auris* treatment is not combination therapy but monotherapy with a single echinocandin, despite the inherent resulting risk of resistance emergence^27^. Alarmingly, the number of echinocandin resistant cases in the United States has tripled in recent years, from six cases before 2020, to 19 cases in 2020 alone^28^.

Echinocandin resistance can emerge through variants in hot-spot sites of the β-glucan synthase gene *FKS1* that affect drug binding (amino acid positions 635-642, 1351-1358, 686-696). Though clades exhibit specific variants^10^, testing 304 global strains from four clades indicated that the most common variant conferring echinocandin resistance was *FKS1* S639P^20^. Multiple viable variants exist, including F635C/I/L/Y/Δ/S/L, S639F/P/T/Y, D642Y/H (hot-spot 1), R1354S/H (hot-spot 2), M690I and W691L/C (hot-spot 3)^29–34^. The presence/absence of a single hot-spot variant can correlate with resistance in clinical case series, such as the S639F variant and resulting echinocandin MICs of ≥2 μg/mL for caspofungin and ≥4 μg/mL for anidulafungin and micafungin^35,36^. *In vitro* experiments demonstrate drug resistance and increased virulence in murine candidiasis with specific variants, such as F635Y^37^ or R1354H^38^. The relative contributions of non-*FKS* mechanisms to multifactorial echinocandin resistance, such as efflux pumps or transcriptional regulators^39^, remain less quantified but may act synergistically with *FKS* target-site mutations. Not all hot-spot variants confer echinocandin resistance in isolation, and no single variant can be used as a species-wide proxy for drug failure using e.g. rapid PCR diagnostic testing^32,40^.

Across four clades, azole resistance has most commonly been attributed to the Y132F variant^20^; K143R^41^ and VF125AL^42^ also confer resistance, confirmed by Cas9-ribonucleoprotein (RNP) mediated transformation^43^ and site-directed mutagenesis and hetorologous expression in *S. cerevisiae* for F126L^44^, Y132F, and K143R^45^. Polyene resistance can occur secondary to clinically rare variations in sterol biosynthesis genes^46^, such as sterol-methyltransferase frame-shift indels in *ERG6*^47^. Clinical flucytosine resistance has occurred with variants in uracil phosphoribosyltransferase *FUR1* (e.g. F211I)^48^ and has emerged in pan-resistant strains with *FUR1* deletion, or variants in flucytosine deaminase *FCY1*^30^. Further important mechanisms of drug resistance include variants in multi-drug transporters such as *CDR1* (V704L)^30^ or their controlling transcriptions factors, such as *TAC1b* (S611P)^49^ and *MRR1* (N647T)^50^.

In this study, we systematically analyse publicly available WGS for *C. auris* up to August 2024 to understand its global genomic epidemiology and describe standing variation related to drug resistance.

## Results

### Global spread tracked by adoption of whole genome sequencing

We analysed 13,200 Illumina paired-end whole genome sequence (WGS) datasets spanning six continents and 27 years (1997 to 2024, **Figure S1**). Of these, 13,193 were downloaded from the NCBI Sequence Read Archive, representing 97 sequencing projects (median 13 isolates per project, range 1-2987, **Table S1**) contributed by 62 clinical, public health, and academic centers across 41 countries. We additionally generated WGS for seven isolates representing clades I-IV from critically unwell patients in a single hospital in Tlemcen, Algeria^14^.

Most samples were from clinical or environmental sources (*n* = 12,920/13,200, 97.9%), with a small minority from laboratory-derived or *in vitro* micro-evolved isolates (*n* = 280/13,200, 2.1%). Clinical and environmental isolates originated from 38 countries across 6 continents, predominantly North America (*n* = 11,388/12,920, 88.1%), followed by Asia (*n* = 452/12,920, 3.5%), Europe (*n* = 436/12,920, 3.4%), Africa (*n* = 357/12,920, 2.8%), South America (*n* = 257/12,920, 2.0%), and Oceania (*n* = 29/12,920, 0.2%). Most of the clinical and environmental datasets were collected from high income countries (12,035/12,920, 93.2%), with substantially fewer from upper-middle-income countries (*n* = 719/12,920, 5.6%) and LMICs (*n* = 165/12,920, 1.3%). No datasets were collected from low-income countries.

Public repository metadata showed substantial limitations for temporal and host niche resolution. Collection dates were recorded to day level precision for only 7,627 of the 13,200 WGS (57.8%), with the remainder reported to the month only (*n* = 3,814/13,200, 28.9%), year only (*n* = 1,513/13,200, 11.5%), or inferable solely from the dataset release date (*n* = 245/13,200, 1.9%, **Figure 2A-B**). Sample site information was missing or ambiguous for 5,798 of the 12,920 clinical or environmental datasets (44.9%). Among datasets with clearly annoted sample sites, the most commonly reported sources were blood (*n* = 1,983/12,920, 10.3%), epithelial or skin swabs (*n* = 1,789, 13.8%), and urine (*n* = 1,649, 12.8%, **Figure 2C**).

### Global nuclear and mitochondrial phylogenies support six-clade structure

We aligned WGS for 13,200 isolates to the nuclear and mitochondrial genome assemblies of the B8441 v3 reference strain. Nuclear alignments exceeding 10 Mb (80.6% of the reference length) were obtained for 12,644 of 13,200 isolates (95.8%), while mitochondrial alignments exceeding 20 kb (70.9% of the reference length) were obtained for 12,510 isolates (94.8%). From these alignments, we identified 506,505 phylogenetically informative sites (4.1% of the nuclear genome assembly length) and 362 mitochondrial sites (1.3% mitochondrial genome assembly length), which were used to infer approximately maximum likelihood phylogenies. Across the nuclear and mitochondrial multiple alignments, only 82 (0.016%) and 15 (0.053%) variant sites, respectively, were covered in all samples. We also detected 77 of 12,644 isolates (0.6%) had identical nuclear sequences, while 10,807 of 12,510 isolates (86.4%) had identical mitochondrial sequences.

Nuclear and mitochondrial phylogenies supported the six-clade structure (**Figure 1A-B**): clade III (*n* = 5,591/12,644, 44.2%), clade I (*n* = 5,275/12,644, 41.7%), clade IV (*n* = 1,683/12,644, 13.3%), clade II (*n* = 81/12,644, 0.6%), clade V (*n* = 6/12,644, <0.1%), and clade VI (*n* = 6/12,644, <0.1%), with two outlier isolates derived from axillary/groin swabs in the USA situated between clade I and IV in the nuclear phylogeny (**Table 1**). Based on available metadata, the earliest WGS for each clade were identified in France (clade I, 2007), Japan (clade II, 1997), Kenya (clade III, 2011), South Africa (clade IV, 2009), Iran (clade V, 2019), and Singapore (clade VI, 2018, **Figure 1A**). The mitochondrial genome alignment of the earliest historical clade IV isolate (South Africa, 2009) did not meet the minimum alignment thresholds (20 kb) for inclusion in the mitochondrial phylogeny **Figure 1B**). We note that the isolation date of the original description of *C. auris* (clade II) from a case of otomycosis in Japan^7^ has been reported as 2005^74^.

**Figure 1:**
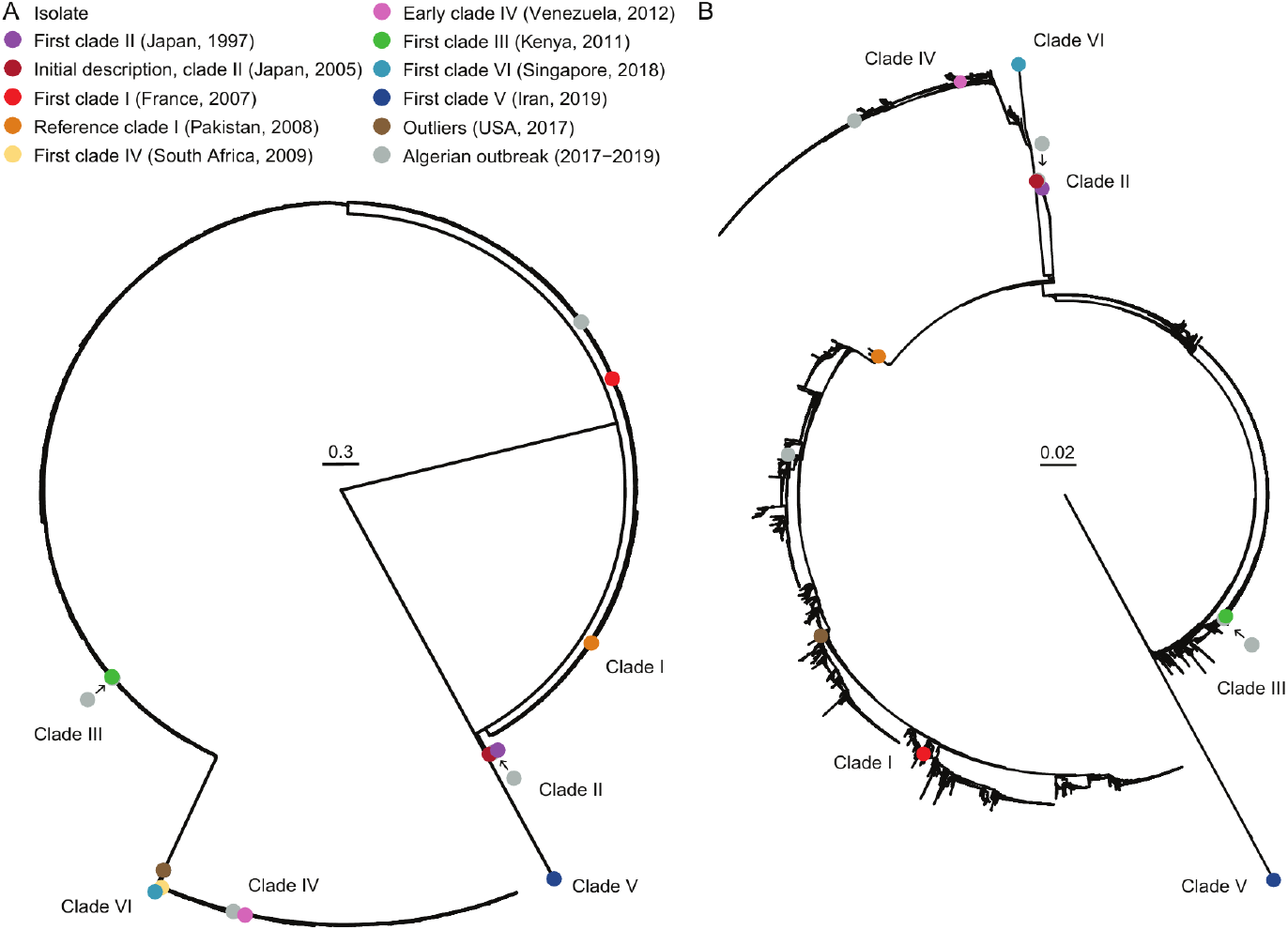
Global genomic population structure: **(A)** Nuclear and **(B)** mitochondrial midpoint rooted phylogenies estimated by approximately maximum likelihood of 506,506 and 362 phylogenetically informative sites across 12,644 and 12,510 isolates (plus reference), respectively. Scale bar indicates substitutions per site.

**Figure 2:**
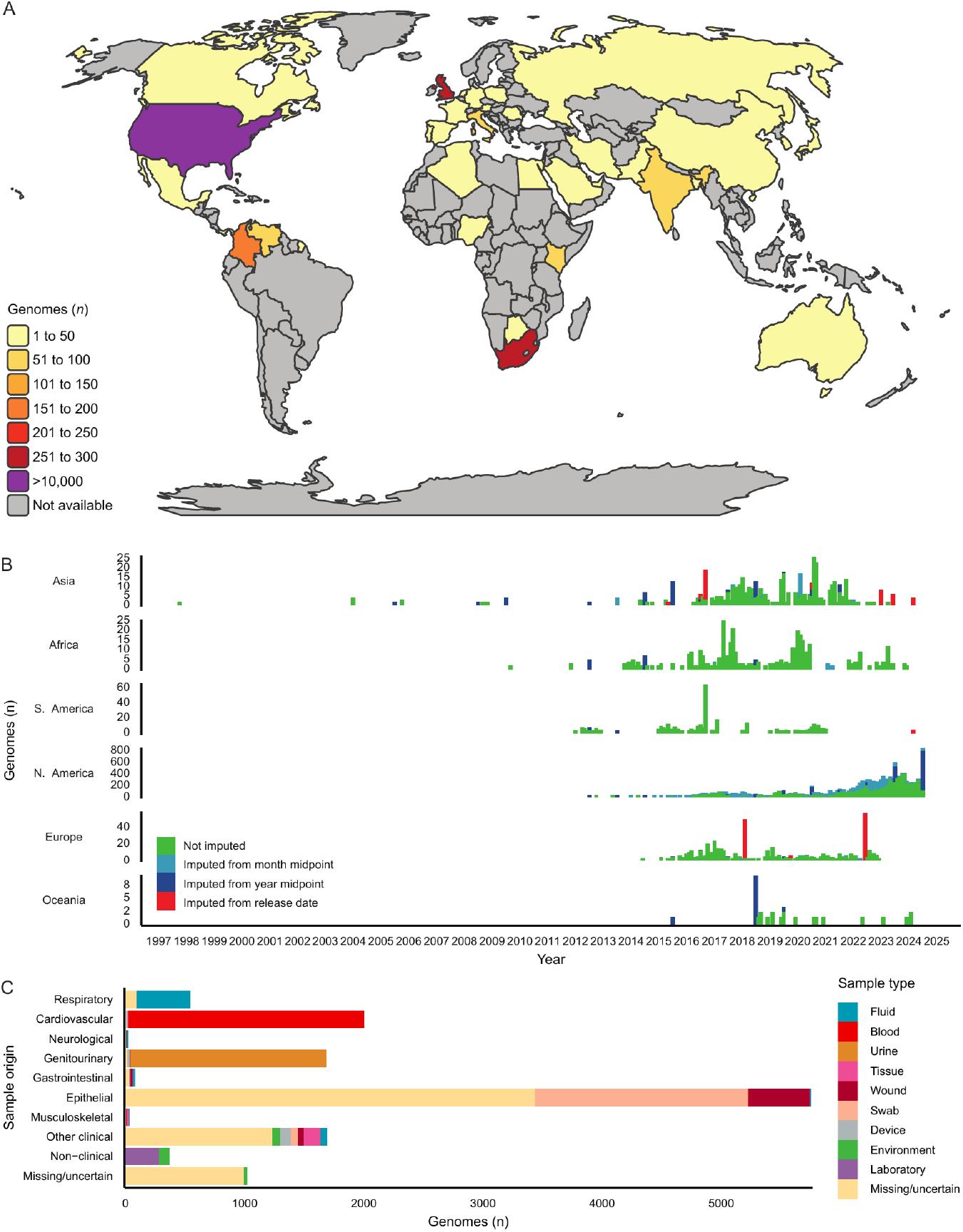
Global genomic epidemiology: **(A)** World map of countries contributing whole genome sequences (WGS) to *C. auris* in this study. **(B)** Epidemiological curve of WGS data across six continents coloured by precise dating source. **(C)** Isolation sources for WGS data, including clinical and non-clinical isolates.

**Table 1:**
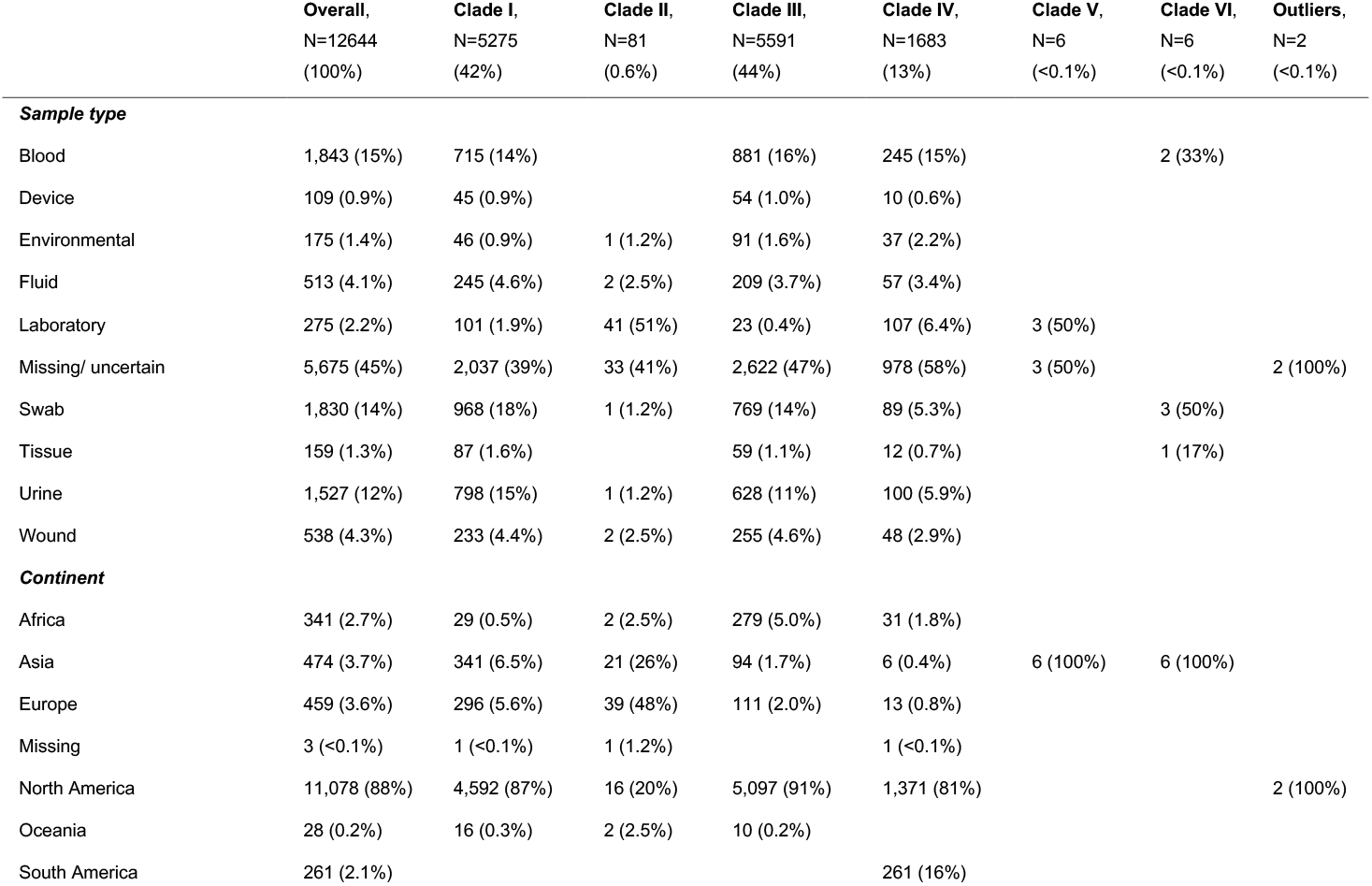
Distribution of estimated clades and sample collection type and continent of collection.

|  | **Overall,**<br>**N=12644**<br>**(100%)** | **Clade I,**<br>**N=5275**<br>**(42%)** | **Clade II,**<br>**N=81**<br>**(0.6%)** | **Clade III,**<br>**N=5591**<br>**(44%)** | **Clade IV,**<br>**N=1683**<br>**(13%)** | **Clade V,**<br>**N=6**<br>**(<0.1%)** | **Clade VI,**<br>**N=6**<br>**(<0.1%)** | **Outliers,**<br>**N=2**<br>**(<0.1%)** |
| --- | --- | --- | --- | --- | --- | --- | --- | --- |
| **Sample type** |  |  |  |  |  |  |  |  |
| Blood | 1,843 (15%) | 715 (14%) |  | 881 (16%) | 245 (15%) |  | 2 (33%) |  |
| Device | 109 (0.9%) | 45 (0.9%) |  | 54 (1.0%) | 10 (0.6%) |  |  |  |
| Environmental | 175 (1.4%) | 46 (0.9%) | 1 (1.2%) | 91 (1.6%) | 37 (2.2%) |  |  |  |
| Fluid | 513 (4.1%) | 245 (4.6%) | 2 (2.5%) | 209 (3.7%) | 57 (3.4%) |  |  |  |
| Laboratory | 275 (2.2%) | 101 (1.9%) | 41 (51%) | 23 (0.4%) | 107 (6.4%) | 3 (50%) |  |  |
| Missing/ uncertain | 5,675 (45%) | 2,037 (39%) | 33 (41%) | 2,622 (47%) | 978 (58%) | 3 (50%) |  | 2 (100%) |
| Swab | 1,830 (14%) | 968 (18%) | 1 (1.2%) | 769 (14%) | 89 (5.3%) |  | 3 (50%) |  |
| Tissue | 159 (1.3%) | 87 (1.6%) |  | 59 (1.1%) | 12 (0.7%) |  | 1 (17%) |  |
| Urine | 1,527 (12%) | 798 (15%) | 1 (1.2%) | 628 (11%) | 100 (5.9%) |  |  |  |
| Wound | 538 (4.3%) | 233 (4.4%) | 2 (2.5%) | 255 (4.6%) | 48 (2.9%) |  |  |  |
| **Continent** |  |  |  |  |  |  |  |  |
| Africa | 341 (2.7%) | 29 (0.5%) | 2 (2.5%) | 279 (5.0%) | 31 (1.8%) |  |  |  |
| Asia | 474 (3.7%) | 341 (6.5%) | 21 (26%) | 94 (1.7%) | 6 (0.4%) | 6 (100%) | 6 (100%) |  |
| Europe | 459 (3.6%) | 296 (5.6%) | 39 (48%) | 111 (2.0%) | 13 (0.8%) |  |  |  |
| Missing | 3 (<0.1%) | 1 (<0.1%) | 1 (1.2%) |  | 1 (<0.1%) |  |  |  |
| North America | 11,078 (88%) | 4,592 (87%) | 16 (20%) | 5,097 (91%) | 1,371 (81%) |  |  | 2 (100%) |
| Oceania | 28 (0.2%) | 16 (0.3%) | 2 (2.5%) | 10 (0.2%) |  |  |  |  |
| South America | 261 (2.1%) |  |  |  | 261 (16%) |  |  |  |

### Genomic plasticity in *C. auris* includes copy number variation in *ERG11*

We detected widespread gene copy number variation (CNV) representing structural variation through gene-level deletion or amplification across the *C. auris* dataset. Our analysis focused on isolates from clinical or environmental origin rather than laboratory or micro-evolved origin (*n* = 12,369, **Figure S2**). Clade-level summaries revealed extensive genome-wide CNV in the most heavily sampled clades (I, III and IV), including large contiguous regions of increased copy number consistent with structural duplications, particularly on Chromosome 1 in clades I and IV. Higher rates of CNV were identified near contig ends consistent with sub-telomeric instability, and whole contig depth of coverage increases suggest natural aneuploidy. For example, genomic signatures of aneuploidy were detected in chromosomes 5 and 6 in clade II. Together, these patterns indicate substantial genomic plasticity across global *C. auris* populations, with gene amplification and deletion likely contributing to adaptive potential.

Among 47 candidate antimicrobial resistance (AMR) genes, we detected the full range of copy number variation, from potential deletion to duplication and amplification (**Figure S3A**). Seven genes demonstrated evidence of quintuple or greater copy number (normalised depth of coverage ≥4.5), including sterol synthesis genes *ERG11_1448* and *ERG10_3730*, multi-drug efflux pumps *SNQ2_4452* and *MDR1_4113*, and transcription factors *FLO8_0401, FLO8_0402*, and *MRR1b_2931* **Figure S3B**). The highest CNV for *ERG11_1448* was identified in isolates derived in India between 2019 and 2020, driven by seven isolates with normalised depth of coverage ranging from 10.2-15.3. Globally, *ERG11_1448* duplications and amplifications were most commonly identified in North America (*n* = 1,834), followed by Africa (*n* = 54), Asia (*n* = 30), South America (*n* = 11), Europe (*n* = 8) and Oceania (*n* = 3, **Figure S3C**).

### Clade-specific loss of function mutations in candidate antimicrobial resistance (AMR)-related genes

We annotated 90,302 unique coding-region variants across 12,641 of 12,644 isolates, representing a total of 81.6 million variants. Among clinical and environmental isolates, 866 distinct non-synonymous, read-through, nonsense and indel variant types were detected across 47 AMR-related genes. In total, 562,861 AMR-related variants were identified across 12,364 of 12,369 epidemiologically-derived isolates, corresponding to a mean of 45.5 potentially impactful AMR-related variants per isolate.

After excluding two outlier samples and one with unresolvable missing continental metadata (*n* = 12,366), we assessed variants predicted to have significant functional impact, such as loss-of-function mutations (**Figure 3**). Several loss-of-function mutations were clade-specific. For example, most clade IV isolates cary a readthrough mutation in *ERG5_1372*, whereas clade II isolates predominantly harbor a frameshift deletion in *ERG5_1372*. Transcription factors *FLO8_0401* and *FLO8_0402* showed high prevalence of frameshift indels in clades II and III and a readthrough mutation in clade I, respectively. In contrast, some key resistance-associated genes, including *ERG11_1448* and *FKS1_0964*, showed an absence of nonsense, readthrough, or frameshift mutations across the global dataset.

**Figure 3:**
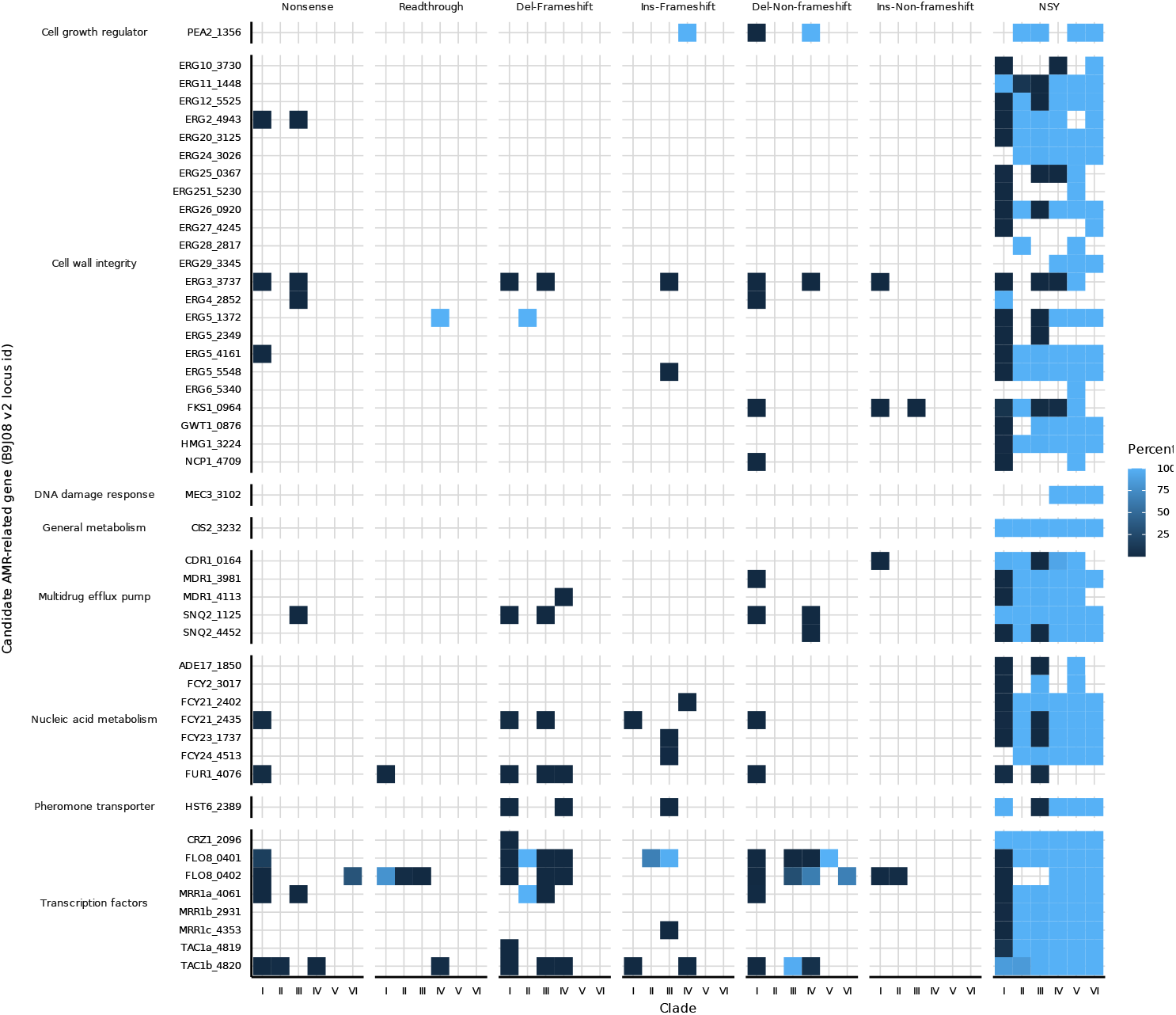
Global survey of significant mutations in epidemiological origin samples. Percentage indicates the percent of isolates per clade containing any number of each type of mutation.

### Expansion of *FKS1* hot-spot diversity

We identified a striking expansion of *FKS1* hot-spot variation relative to existing reports in databases. Where databases have so far recorded a range of approximately eleven different mutations across three hot-spots (amino acid positions 635-642, 1351-1358, 686-696), we identified a total of 36 different mutations, most of which were found in clade I, with a per-clade percentage of 0.02-1.59%. The majority of the 61 clinical or environmental isolates displaying one of 18 emerging *FKS1* hot-spot variants derived from North America (clade I, *n* = 38, clade III, *n* = 14, clade IV, *n* = 5), totalling 57/61 (93.4%). The remainder derived from Africa (clade III, *n* = 2), Asia (clade I, *n* = 1), and Europe (clade I, *n* = 1). The majority of *FKS1* hot-spot mutations were derived within the last 5 years (**Figure 4**), potentially consistent with a real-time emergence of new escape variants. This pattern aligns with concerns in countries where echinocandin monotherapy is used (such as in North America) and may select for the development of resistance. Variants for Algerian isolates are reported in the Supplementary information appendix (**Supplementary results, Table S2**).

**Figure 4:**
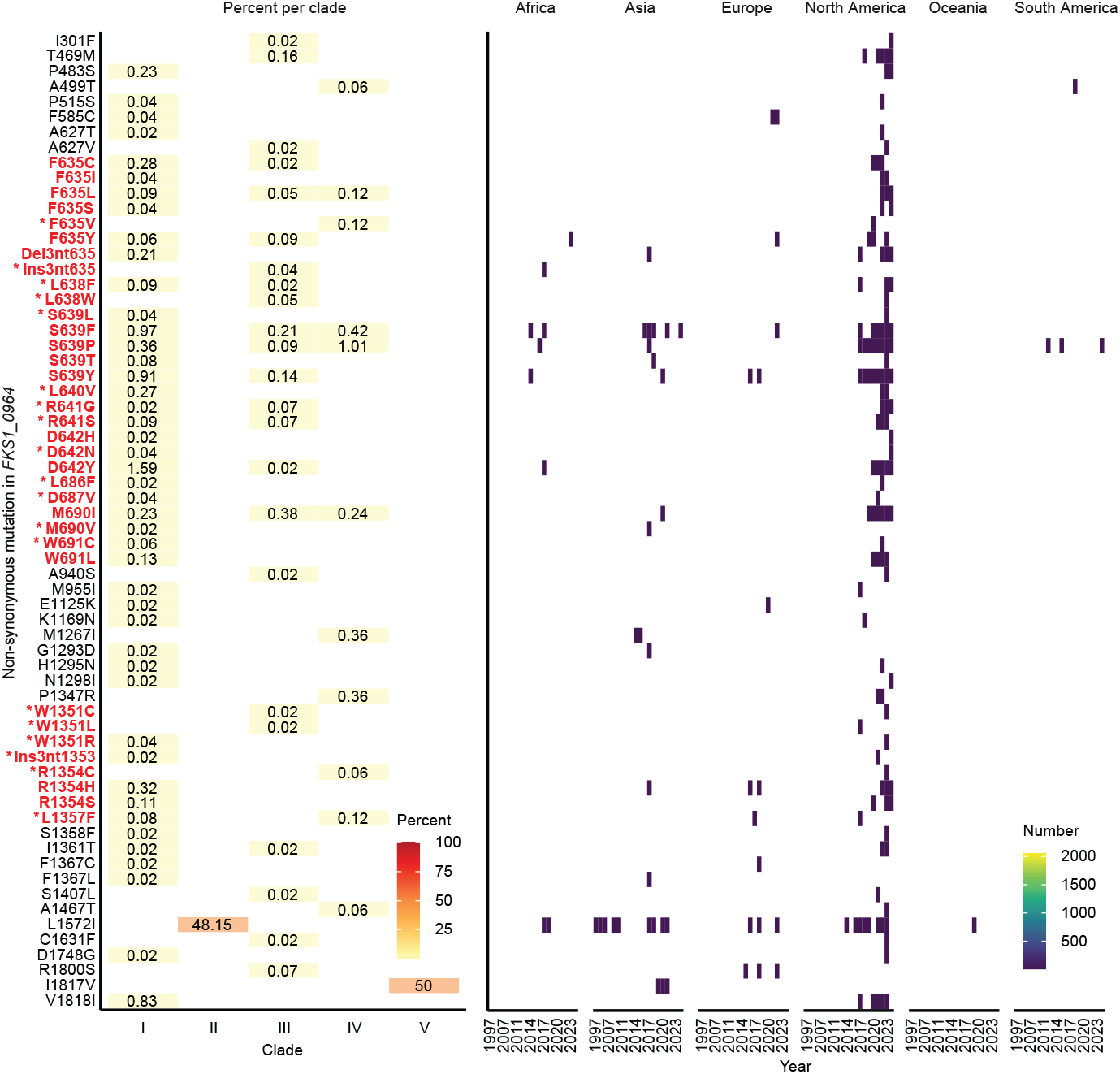
Global survey of *FKS1* mutations in epidemiological origin samples. Left-hand panel indicates the percentage of isolates from each clade exhibiting each non-synonymous mutation. Right-hand panel shows the number of isolates derived from each continental location per year with each corresponding mutation. Mutations in red/bold are in *FKS1* hot-spot regions, with variants not represented in existing databases starred.

To investigate associations between genotype and infection niche, we analysed isolates from blood, urine, and axillary or groin skin swabs within clades I, III, and IV. Blood-derived isolates numbered 715, 881, and 245 for clades I, III, and IV, respectively; urine-derived isolates numbered 798, 628, and 100; and skin-derived isolates numbered 1,209, 1,213, and 851. We performed genome-wide association using per-variant presence or absence matrices with Fisher’s exact tests and Benjamini-Hochberg correction and report the ten most significant variants for each comparison (**Figure S4**).

Across 44,309 variants spanning 5,108 loci, we identified 1,791 variants across 953 loci with an adjusted -log_10_(p-value) of 5.0 or greater. In the clade I comparison of blood *vs* urine, the most significant variant for was *FKS1* S639F, identified in 0.6% of blood-derived isolates and 4.5% of urinary isolates (-log_10_(p-value) 6.33). In clade IV comparison, seven variants in the putative mannosyltransferase *MNN1* were significantly enriched in blood isolates, including T509A, F596S, N664T, F713L, and a readthrough mutation at amino acid position 755, with frequencies of 96.7-98.4% in blood isolates compared with 72-79% in urine isolates. A nonsense mutation in membrane flippase *OSH2* at position 66 was also more common in clade IV blood samples (46.9%) than in urinary samples (34.9%).

Comparisons between blood and skin isolates identified additional clade-specific associations. In clade I, significant differences were observed in known virulence factors, including the adhesin *IFF4* (E570D, 4.2% *vs* 0.8%) and the secreted aspartyl protease *SAP9* (frameshift insertion at amino acid position 273, 7.0% *vs* 18.9%). In clade III, four variants in transcription factor *PRD1* (R193K, N401I, N524D, and P666A) were more common in blood-derived than skin-derived isolates (75.3-75.5% *vs* 42.4-42.55%).

## Discussion

We present a global genomic epidemiology survey of the emerging human fungal pathogen *C. auris*, demonstrating the distribution of six major clades across six continents over nearly three decades (1997-2024). While whole genome sequencing has rapidly expanded in use, its application remains highly uneven, resulting in sampling bias. Nevertheless, the scale and diversity of available global sequencing data enable targeted analyses, including the identification of novel *FKS1* mutations consistent with selection for echinocandin drug resistance, particularly in urinary isolates. These insights highlight the strengths of scale and scope in studying the comprehensive global genomic epidemiology of an emerging pathogen.

Our survey highlights profound global inequalities in sequencing capacity. Over 90% of publicly available sequences originate from high income countries, despite these countries comprising less than 20% of the global population^75^. Addressing this imbalance by improving sequencing infrastructure and data integration in low and lower-middle-income countries is essential for effective genomic surveillance and understanding of pathogen emergence in settings no less affected by disease burden. Failure to include such regions inevitably results in an incomplete representation of species diversity, reinforcing existing biases and limiting epidemiological inference^76^.

We incorporated a small number of isolates from an outbreak in a single ICU in Algeria, where screening of 87 patients revealed the co-circulation of four clades (I-IV)^14^. Algeria experiences high levels of migration^77^, and the phylogenetic placement of Algerian isolates (2017-2019) showed proximity to early representatives of multiple clades. As we have previously argued, assigning geographical identities to the clades is both unhelpful and unethical^16^, particularly in the context of uneven surveillance and globalised human movement. Inclusion of isolates from under-represented regions in global genomic epidemiology is essential, but limited numbers in this study restrict the strength of conclusions toward inferring transmission routes. Employing more equitable sequencing and using One Health ecological approaches should help overcome such barriers to understanding the evolution of resistance-associated variants.

The conserved nuclear clade structure observed globally may reflect several non-exclusive processes. These include a limited number of successful emergent lineages from ancestral reservoir(s), persistent circulation of dominant strains in heavily sampled high-income regions, and the absence of coordinated one-health approaches to recover *C. auris* from environmental niches. These interpretations are further constrained by limitations of public data repositories, where metadata are frequently incomplete, inconsistent, or ambiguous. We recommend improved data standards that require structured metadata alongside genome submission, including accurate isolation dates, geographic location at city level, and clinical body site. Without such information, sequencing risks becoming an exercise in data accumulation rather than meaningful biological or epidemiological insight.

We discovered a substantial expansion of *FKS1* hot-spot variation. Mutations in *FKS1* hot-spots can alter echinocandin binding to the β-1,3-glucan synthase target site and are a well-established mechanism of drug resistance across yeast species^78^. Prior surveys of *C. auris* identified approximately eleven hot-spot mutations across three regions of *FKS1*: F635L/Y/Δ, S639F/P/T/Y, D642Y (hot-spot 1), R1354S/H (hot-spot 2), and W691L (hot-spot 3)^31^. Our global analysis expands this catalogue by adding an additional 24 hot-spot variants, many of which have not been previously reported. Only six of these mutations have been identified elsewhere (F635C^30^, F635I^33^, F635S/L and D642H^32^, M690I^29^, and W691C^34^, indicating that there are likely to be at least 18 newly observed variants of potential clinical relevance. We propose the development of integrated and openly reported phenotypic surveillance (including antifungal MIC values) alongside genomic surveillance AMR-related databases to integrate phenotypic data as a highly effective approach for monitoring the impact of evolution in this priority pathogen.

Emerging *FKS1* mutations concentrated in recent North American isolates, where echinocandin monotherapy is available and widely used as first-line treatment^8^. The major factor contributing to this signal is likely to be the excessive variability in genomic surveillance intensity: 11,078 (88%) of surveyed isolates derived from N. America. Other high-income countries are likely to follow international consensus on echinocandin monotherapy, and lower-income countries may suffer from lack of access to echinocandin antifungals, but our survey does not describe extensive novel *FKS1* hot-spot variants in these setting. The emergence of so many *FKS1* hot-spot variants is consistent with drug-driven selection generating a diverse spectrum of resistance alleles. The breadth of *FKS1* variants also underscores the limitation of real-time diagnostic PCR assays^40^, which cannot detect novel hot-spot variants and are therefore unsuitable for comprehensive resistance surveillance or clade differentiation. Even when focused on a single locus, our results demonstrate the value of large-scale genomic epidemiology to uncover potential clinically significant variation that has likely been underestimated to date. Nevertheless, the absence of quantitative clinical data (such as MIC values) and limitations around metadata completeness call for a cautious interpretation of the clinical impact of newly identified variants. Targeted antifungal susceptibility testing for isolates with emerging variation in key AMR-related loci could provide better real-time detection of novel variants of major impact.

We examined genome plasticity through CNV inferred from measurement of normalised depth of coverage. We detected up to 15 copies of *ERG11* in isolates from India collected between 2019 and 2020. While *ERG11* amplification has been reported in laboratory-aged isolates and linked to azole resistance and therapeautic failure in *Galleria mellonella* infection^79^, we are unaware of prior reports documenting such amplification in clinical isolates. Genome-wide CNV was most apparent in clades I, III and IV, likely reflecting biological variation and increased sampling depth. However, CNV inference from short-read data remains sensitive to technical factors such as library preparation, and confirmation using assembly-based approaches, ideally incorporating long-read sequencing, is necessary. Such approaches have previously demonstrated large sub-telomeric amplifications, including ten-fold expansion of adhesin *ALS4* in urinary isolates^80^. Sub-telomeric instability, often driven by ectopic recombination^81^, is a plausible mechanism for adaptive gene gain and loss, and *Candida* species are enriched for adhesins in these regions^82^. In *C. auris*, sub-telomeric adhesins *ALS4* and *SCF1* contribute to virulence^80,83^ and are up-regulated during *in vivo* murine catheter infection^84^. Further exploratory analysis could assess the functional enrichment of gene gain/loss at telomeres in the global data, complemented by assembly-based methods informed by long read sequencing^85^ with *in vitro* antifungal susceptibility testing of isolates carrying CNVs to document the penetrance and clinical impact of genomic innovation.

Using a conservative GWAS approach, we identified genotypic variation associated with clinical body site infection niche. The most significant variant enriched in urinary isolates relative to bloodstream isolates in clade I was *FKS1* S639F. This mutation is a known echinocandin resistance determinant that can emerge during therapy and lead to treatment failure^86,87^. Echinocandins achieve poor urinary concentrations, potentially creating conditions conducive to resistance emergence in this niche^88^. Additional niche-associated variants were identified in known virulence factors, including a frameshift mutation in secreted protease *SAP9* in the skin niche compared to bloodstream isolates in clade I^89^. These findings suggest that the colonisation of multiple anatomical sites exposes *C. auris* populations to distinct selective pressures, facilitating ongoing adaptation that may compromise treatment efficacy and contribute to adverse clinical outcomes.

This study demonstrates the power of global genomic epidemiology to resolve population structure, identify emerging resistance-associated variants, and detect genotype-phenotype associations with clinical relevance. By systematically curating public data and analysing 12,644 nuclear genomes, we confirmed a six-clade population structure and document substantial geographic sampling bias. Our results reveal extensive previously unrecognised *FKS1* diversity and show how large-scale datasets can inform statistical genetic analyses with implications for antifungal resistance, public health, and policy. We outline the need for targeted phenotypic integration to translate genomic surveillance into meaningful clinical impact and evolutionary insight. Alongside humanitarian and One Health imperatives, these findings demonstrate the need to study emerging fungal pathogens within their global context.

## Methods

### Search strategy

The National Center for Biotechnology Information Sequence Read Archive (NCBI SRA) was searched for Illumina paired end whole genome sequencing (WGS) reads for “*Candida auris*” on 23rd August 2024. The Run Selector function was used to obtain metadata and sequence accession lists^51,52^. Metadata were examined to elucidate geographical and clinical isolation sources and collection dates where available, or assigned to mid-point of month/year where only this was given, or given an imputed value where public release date was equal to sample collection date to adjust for missing data (**Supplementary methods**). Clinical samples were assigned to type (patient, device, object or environment) organ system (cardiovascular, respiratory, epithelial *etc*.), anatomical location, and sample type (swab, central venous catheter, tissue, *etc*.). Additional country and source details were derived manually from additional data where possible, including primary literature (**Table S1**) and primary project metadata provided directly by article authors^15,22^. Country income status was derived from the latest available World Bank data^53^. Maps were plotted with tmaptools v.3.1-1 and tmap v.3.99.9002^54^.

### Genome sequencing and variant calling

Additional strains from the Algerian outbreak (*n* = 7)^14^ were plated from microdiscs onto YPD agar (Sigma-Aldrich, UK), stored in YPD broth with 25% glycerol at -80 °C, inoculated into 5 mL overnight in a shaking incubator at 30 °C, 200 rpm, and extracted using the DNeasy Plant Pro kit (QIAgen, #69204) according to manufacturer’s instructions. DNA samples were quantified using the Qubit 4.0 fluorometer and integrity checked on agarose gel. Library preparation was performed with the NEB Ultra II FS DNA Low Volume kit (Band ILL7) and sequencing was performed by the Exeter Sequencing Service on the Illumina NovaSeq SP 300 (Band ISN3).

For publicly available genomes, SRA FASTQ files were downloaded using SRA tools v.3.1.0 fastq-dump and checked for quality using FastQC v.0.12.1^55^ and MultiQC v.1.21^56^. After indexing with SAMtools v.1.17^57^, alignment was performed using bwa v.0.7.17 with soft clipping for supplementary alignments and marking of shorter split hits as secondary^59^ using the B8441 v.3 (GCA 002759435) nuclear reference^10,60^. Variant calling was performed with the snippy v.4.6.0 pipeline^61^, including a maximum clip length of 10, SAMtools sort by read name, SAMtools add score to read pair with fixmate, and SAMtools duplicate read removal. For variant calling, we used FreeBayes-parallel v.1.3.6^62^ with a ploidy level of 2, a minimum allele count of 2, a minimum alternate allele fraction of 0.05, a minimum coverage of 10, a minimum of 1 bit per base Shannon entropy to detect interrupted repeats, minimum Phred-scaled base quality of 13, minimum mapping quality of 60. Variants were filtered with bcftools v.1.17^63^ view for a quality score >100, a minimum depth of 10, a minimum allele observation count fraction of depth of 0. We normalised to the reference with vt v.2015.11.10^64^ and used bcftools to compress and index the vcf.

After alignment, we used SAMtools pileup to calculate depth and breadth of coverage at each position, informing depth of coverage (DOC) normalised to mean DOC across the genome for non-overlapping 10 kb sliding windows or for each gene locus. To perform a screening GWAS, we made use of isolates with verifiable epidemiological origin in blood, urine, or axillary/groin skin. Two-tailed Fisher’s exact tests were performed using per-variant two-by-two presence/absence matrices with Benjamini-Hochberg testing for multiple correction across each comparison using R v.4.4.0 (stats::fisher.test, stats::p.adjust).

### Phylogenomic analysis

Core multi-fasta alignments were created with snippy-core v.4.6.0^61^, filtering for a minimum coverage of 10 Mb (nuclear) or 20 kb (mitochondrial) in batches of 1000 until a successful core phylogeny was identified for each batch. Following this, we then determined phylogenetically informative sites using snp-sites v.2.5.1^65^ across all isolates with adequate coverage as above. We calculated approximately maximum likelihood phylogenies for nuclear and mitochondrial genomes separately VeryFastTree with the GTR rate substitution model^66,67^. Midpoint-rooted trees were plotted with phangorn v.2.12.1^68^, ggtree v.3.14.0^69^, treeio v.1.8.0^70^, ggtreeExtra v.1.14.0^71^ and TDbook v.0.0.6^72^. Differences in numbers of variants between WGS were calculated with snp-dists v.0.8.2^73^.

## Data availability

Raw reads have been deposited on NCBI SRA accession number PRJNA1372636.

## Author contributions

**HG** - analysis, writing - original draft, conceptualisation; **NH** - resources, editing; **HZ** - resources, editing; **DW** - analysis, editing, supervision, conceptualisation; **JR** - analysis, editing, supervision, conceptualisation; **RF** - analysis, editing, supervision, conceptualisation.

## Acknowledgements

We acknowledge funding from the MRC Centre for Medical Mycology at the University of Exeter (MR/N006364/2, MR/V033417/1) and the MRC Doctoral Training Grant MR/P501955/2, Wellcome Trust Career Development Award (215239/Z/19/Z) and Wellcome Trust fellowship (219551/Z/19/Z), and the NIHR Exeter Biomedical Research Centre. The views expressed are those of the authors and not necessarily those of the NIHR or the Department of Health and Social Care. We are grateful to Fadi Bittar for assisting in communication. We also thank the Exeter Sequencing Service facility and support from Wellcome Trust Institutional Strategic Support Fund (WT097835MF), Wellcome Trust Multi User Equipment Awards (WT101650MA and 218247/Z/19/Z), Medical Research Council Clinical Infrastructure Funding (MR/M008924/1) and BBSRC LOLA award (BB/K003240/1), as well as the University of Exeter High-Performance Computing (HPC) facility, funded by the UK MRC Clinical Research Infrastructure Initiative (award number MR/M008924/1).

## Notes

### Competing Interest Statement

The authors have declared no competing interest.

## References

1. Liu, F. et al. Phylogenomic analysis of the Candida auris-Candida haemuli clade and related taxa in the Metschnikowiaceae, and proposal of thirteen new genera, fifty-five new combinations and nine new species. Persoonia - Molecular Phylogeny and Evolution of Fungi 52, 22–43 (2024).

2. Zhang, S. X. et al. Reaffirming the importance of nomenclature stability for Candida auris and its associated disease of candidiasis. Journal of Clinical Microbiology (2025) doi:10.1128/jcm.01550-25.

3. Meis, J. F. & Chowdhary, A. Candida auris: A global fungal public health threat. The Lancet Infectious Diseases 18, 1298–1299 (2018).

4. WHO. World Health Organization (WHO) fungal priority pathogens list to guide research, development and public health action. (2022) Available at: https://www.who.int/publications/i/item/9789240060241.

5. CDC. Centers for Disease Control and Prevention (U.S.): Antibiotic resistance threats in the United States, 2019. (2019) doi:10.15620/cdc:82532.

6. Kim, H. Y. et al. Candida auris - a systematic review to inform the world health organization fungal priority pathogens list. Medical Mycology 62, myae042 (2024).

7. Satoh, K. et al. Candida auris sp. Nov., a novel ascomycetous yeast isolated from the external ear canal of an inpatient in a Japanese hospital. Microbiology and Immunology 53, 41–44 (2009).

8. Lionakis, M. S. & Chowdhary, A. Candida auris infections. New England Journal of Medicine 391, 1924–1935 (2024).

9. Chen, J. et al. Is the superbug fungus really so scary? A systematic review and meta-analysis of global epidemiology and mortality of Candida auris. BMC Infectious Diseases 20, 827 (2020).

10. Lockhart, S. R. et al. Simultaneous emergence of multidrug-resistant Candida auris on 3 continents confirmed by whole-genome sequencing and epidemiological analyses. Clinical Infectious Diseases 64, 134–140 (2017).

11. Suphavilai, C. et al. Detection and characterisation of a sixth Candida auris clade in Singapore: A genomic and phenotypic study. Lancet Microbe 5, (2024).

12. Chen, Z., Lemey, P. & Yu, H. Approaches and challenges to inferring the geographical source of infectious disease outbreaks using genomic data. The Lancet Microbe 5, e81–e92 (2024).

13. Aanensen, D. M. et al. The genomic surveillance gap: Averting the antimicrobial resistance pandemic requires global equity and action. The Lancet Infectious Diseases 26, 122–123 (2025).

14. Zerrouki, H. et al. Emergence of Candida auris in intensive care units in Algeria. Mycoses e13470 (2022) doi:10.1111/myc.13470.

15. Naicker, S. D. et al. Clade distribution of Candida auris in South Africa using whole genome sequencing of clinical and environmental isolates. Emerging Microbes & Infections 10, 1300– 1308 (2021).

16. Gifford, H., Rhodes, J. & Farrer, R. A. The diverse genomes of Candida auris. The Lancet Microbe 5, (2024).

17. Osaigbovo, I. I. et al. Candida auris : A systematic review of a globally emerging fungal pathogen in Africa. Open Forum Infectious Diseases 11, ofad681 (2024).

18. Ibe, C. & Pohl, C. H. Epidemiology and drug resistance among Candida pathogens in Africa: Candida auris could now be leading the pack. The Lancet Microbe 100996 (2024) doi:10.1016/j.lanmic.2024.100996.

19. Yerbanga, I. W. et al. Epidemiology, clinical features and antifungal resistance profile of Candida auris in Africa: Systematic review. Journal of Biosciences and Medicines 12, 126–149 (2024).

20. Chow, N. A. et al. Tracing the evolutionary history and global expansion of Candida auris using population genomic analyses. mBio 11, 15 (2020).

21. Kekana, D. et al. Candida auris clinical isolates associated with outbreak in neonatal unit of tertiary academic hospital, South Africa. Emerging Infectious Diseases 29, 2044–2053 (2023).

22. Kappel, D. et al. Genomic epidemiology describes introduction and outbreaks of antifungal drug-resistant Candida auris. npj Antimicrobials and Resistance 2, 1–10 (2024).

23. Nash, A. et al. MARDy: Mycology Antifungal Resistance Database. Bioinformatics 34, 3233–3234 (2018).

24. Jain, A., Singhal, N. & Kumar, M. AFRbase: A database of protein mutations responsible for antifungal resistance. Bioinformatics 39, btad677 (2023).

25. Bédard, C. et al. FungAMR: A comprehensive database for investigating fungal mutations associated with antimicrobial resistance. Nature Microbiology 10, 2338–2352 (2025).

26. Kumar, C., Arshi, A., Yadav, A., Chavan, P. & Idicula-Thomas, S. CanDRes: Exploring the Mutation Landscape of Candida and its Role in Antifungal Resistance. Infection 54, 509–514 (2025).

27. Wake, R. M. et al. Optimizing the treatment of invasive candidiasis — a case for combination therapy. Open Forum Infectious Diseases 11, ofae072 (2024).

28. Lyman, M. et al. Worsening spread of Candida auris in the United States, 2019 to 2021. Annals of Internal Medicine 176, 489–495 (2023).

29. Carolus, H. et al. Genome-wide analysis of experimentally evolved Candida auris reveals multiple novel mechanisms of multidrug resistance. mBio 12, e03333–20 (2021).

30. Jacobs, S. E. et al. Candida auris pan-drug-resistant to four classes of antifungal agents. Antimicrobial Agents and Chemotherapy 66, e0005322 (2022).

31. Kordalewska, M. et al. Novel non-hot spot modification in Fks1 of Candida auris confers echinocandin resistance. Antimicrobial Agents and Chemotherapy 67, e00423–23 (2023).

32. Zhu, Y. et al. Development and validation of TaqMan chemistry probe-based rapid assay for the detection of echinocandin-resistance in Candida auris. Journal of Clinical Microbiology 61, e01767–22 (2023).

33. Zhang, Y. et al. Two outbreaks and sporadic occurrences of Candida auris from one hospital in China: An epidemiological, genomic retrospective study. Infection 53, 349–358 (2025).

34. Misas, E. et al. A benchmark dataset for validating FKS1 mutations in Candida auris. Microbiology Spectrum 13, e03147–24 (2025).

35. Kordalewska, M. et al. Understanding echinocandin resistance in the emerging pathogen Candida auris. Antimicrobial Agents and Chemotherapy 62, e00238–18 (2018).

36. Jenull, S. et al. Transcriptomics and phenotyping define genetic signatures associated with echinocandin resistance in Candida auris. mBio 13, e00799–22 (2022).

37. Sharma, D. et al. Impact of FKS1 genotype on echinocandin in-vitro susceptibility in Candida auris and in-vivo response in a murine model of infection. Antimicrobial Agents and Chemotherapy 66, e01652–21 (2021).

38. Kiyohara, M. et al. Evaluation of a novel FKS1 R1354H mutation associated with caspofungin resistance in Candida auris using the CRISPR-Cas9 system. Journal of Fungi 9, 529 (2023).

39. Gifford, H. et al. FKS1/2-variant independent mechanisms underlying the emergence of resistance in echinocandin-refractory Candida auris infections. 2026.01.17.700071 (2026) doi:10.64898/2026.01.17.700071.

40. Hou, X. et al. Rapid detection of ERG11-associated azole resistance and FKS-associated echinocandin resistance in Candida auris. Antimicrobial Agents and Chemotherapy 63, e01811–18 (2019).

41. Kwon, Y. J. et al. Candida auris clinical isolates from South Korea: Identification, antifungal susceptibility, and genotyping. Journal of Clinical Microbiology 57, e01624–18 (2019).

42. Tian, S. et al. Genomic epidemiology of Candida auris in a general hospital in Shenyang, China: A three-year surveillance study. Emerging Microbes & Infections 10, 1088–1096 (2021).

43. Rybak, J. M. et al. Delineation of the direct contribution of Candida auris ERG11 mutations to clinical triazole resistance. Microbiology Spectrum 9, e01585–21 (2021).

44. Williamson, B. et al. Impact of Erg11 amino acid substitutions identified in Candida auris clade III isolates on triazole drug susceptibility. Antimicrobial Agents and Chemotherapy 66, e01624–21 (2021).

45. Healey, K. R. et al. Limited ERG11 mutations identified in isolates of Candida auris directly contribute to reduced azole susceptibility. Antimicrobial Agents and Chemotherapy 62, e01427– 18 (2018).

46. Carolus, H. et al. Acquired amphotericin B resistance leads to fitness trade-offs that can be mitigated by compensatory evolution in Candida auris. Nature Microbiology 9, 3304–3320 (2024).

47. Rybak, J. M. et al. In vivo emergence of high-level resistance during treatment reveals the first identified mechanism of amphotericin B resistance in Candida auris. Clinical Microbiology and Infection 28, 838–843 (2022).

48. Rhodes, J. et al. Genomic epidemiology of the UK outbreak of the emerging human fungal pathogen Candida auris. Emerging Microbes & Infections 7, 1–12 (2018).

49. Li, J. et al. Novel ERG11 and TAC1b mutations associated with azole resistance in Candida auris. Antimicrobial Agents and Chemotherapy 65, e02663–20 (2021).

50. Li, J., Coste, A. T., Bachmann, D., Sanglard, D. & Lamoth, F. Deciphering the Mrr1/Mdr1 pathway in azole resistance of Candida auris. Antimicrobial Agents and Chemotherapy 66, e00067–22 (2022).

51. Wheeler, D. L. et al. Database resources of the National Center for Biotechnology Information. Nucleic Acids Research 36, D13–D21 (2008).

52. Shumway, M., Cochrane, G. & Sugawara, H. Archiving next generation sequencing data. Nucleic Acids Research 38, D870–871 (2010).

53. World Bank. World Bank Country and Lending Groups (2024). Available at: https://datahelpdesk.worldbank.org/knowledgebase/articles/906519-world-bank-country-and-lending-groups

54. Tennekes, M. Tmap: Thematic maps in R. Journal of Statistical Software 84, (2018).

55. Andrews, S. FastQC: A quality control tool for high throughput sequence data. Babraham Bioinformatics (2010).

56. Ewels, P., Magnusson, M., Lundin, S. & Käller, M. MultiQC: Summarize analysis results for multiple tools and samples in a single report. Bioinformatics 32, 3047–3048 (2016).

57. Li, H. et al. The Sequence Alignment/Map format and SAMtools. Bioinformatics 25, 2078–2079 (2009).

58. Burrows, M. & Wheeler, D. J. A block-sorting lossless data compression algorithm. Systems Research Center Research Report 124, (1994).

59. Li, H. & Durbin, R. Fast and accurate short read alignment with Burrows–Wheeler transform. Bioinformatics 25, 1754–1760 (2009).

60. Cauldron, N. C., Shea, T. & Cuomo, C. A. Improved genome assembly of Candida auris strain B8441 and annotation of B11205. Microbiology Resource Announcements 3, e00512–24 (2024).

61. Seemann, T. Snippy: Fast bacterial variant calling from NGS reads. (2015). Available at: https://github.com/tseemann/snippy

62. Garrison, E. & Marth, G. Haplotype-based variant detection from short-read sequencing. (2012) doi:10.48550/arXiv.1207.3907.

63. Danecek, P. et al. Twelve years of SAMtools and BCFtools. GigaScience 10, giab008 (2021).

64. Tan, A., Abecasis, G. R. & Kang, H.M. Unified representation of genetic variants. Bioinformatics (Oxford, England) 31, 2202–2204 (2015).

65. Page, A. J. et al. SNP-sites: Rapid efficient extraction of SNPs from multi-FASTA alignments. Microbial Genomics 2, e000056 (2016).

66. Price, M. N., Dehal, P. S. & Arkin, A. P. FastTree 2 – approximately maximum-likelihood trees for large alignments. PLOS ONE 5, e9490 (2010).

67. Piñeiro, C., Abuín, J. M. & Pichel, J. C. Very Fast Tree: Speeding up the estimation of phylogenies for large alignments through parallelization and vectorization strategies. Bioinformatics 36, 4658–4659 (2020).

68. Schliep, K. P. Phangorn: Phylogenetic analysis in R. Bioinformatics 27, 592–593 (2011).

69. Yu, G., Smith, D. K., Zhu, H., Guan, Y. & Lam, T. T.-Y. Ggtree: An R package for visualization and annotation of phylogenetic trees with their covariates and other associated data. Methods in Ecology and Evolution 8, 28–36 (2017).

70. Wang, L.-G. et al. Treeio: An R package for phylogenetic tree input and output with richly annotated and associated data. Molecular Biology and Evolution 37, 599–603 (2020).

71. Xu, S. et al. ggtreeExtra: Compact visualization of richly annotated phylogenetic data. Molecular Biology and Evolution 38, 4039–4042 (2021).

72. Yu, G. Data Integration, Manipulation and Visualization of Phylogenetic Trees. (2022).

73. Seemann, T. Tseemann/snp-dists. (2024). Available at: https://github.com/tseemann/snp-dists

74. Sekizuka, T. et al. Clade II Candida auris possess genomic structural variations related to an ancestral strain. PLOS ONE 14, e0223433 (2019).

75. Fantom, N. & Serajuddin, U. The World Bank’s classification of countries by income. World Bank Group Development Economics Data Group 7528, (2016).

76. Osaigbovo, I. I. et al. The Nairobi Declaration 2023: A commitment to address deadly yet neglected fungal diseases in Africa. Medical Mycology 62, myad141 (2024).

77. Groenewold, W.G.F. (George)., de Beer, J.A.A. (Joop). & de Valk, H.A.G. (Helga). Prospects of labour migration pressure in Algeria, Morocco, Tunisia and Turkey. Genus 72, 8 (2016).

78. Johnson, M. E. & Edlind, T. D. Topological and mutational analysis of Saccharomyces cerevisiae Fks1. Eukaryotic Cell 11, 952–960 (2012).

79. Bhattacharya, S., Holowka, T., Orner, E. P. & Fries, B. C. Gene duplication associated with increased fluconazole tolerance in Candida auris cells of advanced generational age. Scientific Reports 9, 5052 (2019).

80. Bing, J. et al. Clinical isolates of Candida auris with enhanced adherence and biofilm formation due to genomic amplification of ALS4. PLOS Pathogens 19, e1011239 (2023).

81. Anderson, M. Z., Wigen, L. J., Burrack, L. S. & Berman, J. Real-time evolution of a subtelomeric gene family in Candida albicans. Genetics 200, 907–919 (2015).

82. Smoak, R. A., Snyder, L. F., Fassler, J. S. & He, B. Z. Parallel expansion and divergence of an adhesin family in pathogenic yeasts. Genetics 223, iyad024 (2023).

83. Santana, D. J. et al. A Candida auris-specific adhesin, Scf1, governs surface association, colonization, and virulence. Science 381, 1461–1467 (2023).

84. Wang, T. W. et al. Functional redundancy in Candida auris cell surface adhesins crucial for cell-cell interaction and aggregation. Nature Communications 15, 9212 (2024).

85. De Coster, W. & Van Broeckhoven, C. Newest methods for detecting structural variations. Trends in Biotechnology 37, 973–982 (2019).

86. Hirayama, T. et al. Echinocandin resistance in Candida auris occurs in the murine gastrointestinal tract due to FKS1 mutations. Antimicrobial Agents and Chemotherapy 67, e01243–22 (2023).

87. Spruijtenburg, B. et al. Whole genome sequencing analysis demonstrates therapy-induced echinocandin resistance in Candida auris isolates. Mycoses 66, 1079–1086 (2023).

88. Rkieh, L. et al. Outcomes of caspofungin use in the treatment of Candida-related urinary tract infections, a case series. IDCases 28, e01510 (2022).

89. Kim, J.-S.Lee, K.-T. & Bahn, Y.-S. Secreted aspartyl protease 3 regulated by the Ras/cAMP/PKA pathway promotes the virulence of Candida auris. Frontiers in Cellular and Infection Microbiology 13, 1257897 (2023).

