## Supplementary information appendix for "Global genomic epidemiology of *Candida auris*: analysis of 12,644 whole genome sequences from 1997-2024"

Hugh Gifford[a], Nicolas Helmstetter[a], Hanane Zerrouki[b,c,d], Duncan Wilson[a], Johanna Rhodes[e,f], Rhys A. Farrer*[a]

[a]: MRC Center for Medical Mycology, University of Exeter, Geoffrey Pope Building, Stocker Road, Exeter, United Kingdom, EX4 4QD
[b]: Aix-Marseille Université, IRD, APHM, MEPHI, Marseille, France
[c]: IHU-Méditerranée Infection, Marseille, France
[d]: Laboratoire de Microbiologie Appliquée à l'Agroalimentaire, au Biomédical et à l'Environnement, Université de Tlemcen, Tlemcen, Algeria
[e]: School of Biosciences, University of Birmingham, Birmingham, United Kingdom, B15 2TT
[f]: MRC Centre for Global Infectious Disease Analysis, Imperial College London South Kensington Campus, London, United Kingdom, SW7 2AZ
Keywords: Human Fungal Pathogens, Global Genomic Epidemiology, *Candida auris*, Antifungal Resistance, Antimicrobial Resistance, Phylogenomics, Copy Number Variation, Emerging Infectious Pathogens

### Supplementary methods

**Metadata imputation:** The following data columns were informative, for geography: `geo_loc_name_country_continent` and `geo_loc_name_country`; for patient isolation source: `isolation_source`; for date of collection: `Collection_Date`. We parsed dates using the "Collection_Date" column, filtering out all those containing "not collected", "not applicable" or "missing" values (*n* = 296, 2.2%). When there were two dates separated by a "/" we used the first date. A majority of isolates were fully dated (*n* = 7506, 56.9%). For isolates with only year (*n* = 1577, 12.0%) or month (*n* = 3814, 28.9%), we assigned assumed dates at a generic year (July 2$^{nd}$) or month (15$^{th}$) midpoint. For isolates with specified release date and collection date, the mean time between collection and genomic release was 384.9 days (excluding a single run SRR29142435 with a reported collection date in the year 1934), so we imputed a collection date 385 days previous to NCBI SRA release for isolates without collection dates. We also applied this imputation to run SRR29142435. We identified 569 separate terms for different swab origins and assigned categories by type (patient, device, object or environment) organ system (cardiovascular, respiratory, epithelial *etc.*), anatomical location, and sample type (swab, central venous catheter, tissue, *etc.*). Additional country and source details were derived manually from additional data where possible, including primary literature (**Table S1**) and primary project metadata provided directly by article authors[1,2].

**Supplementary results**

We tabulated the seven Algerian isolates, where we did identify a discrepancy in clade identification between our method and the short tandem repeat method used previously for one isolate[3], which could be explained by mixed colonisation, human error, or other causes. We note the presence of the "typical" L1572I non-synonymous mutation in *FKS1* in the two clade II isolates, and similarly common Y132F in *ERG11* in the clade I isolate, as well as several co-inherited *ERG11* mutations in the two clade IV isolates (**Table S2**).

**Supplementary figures**

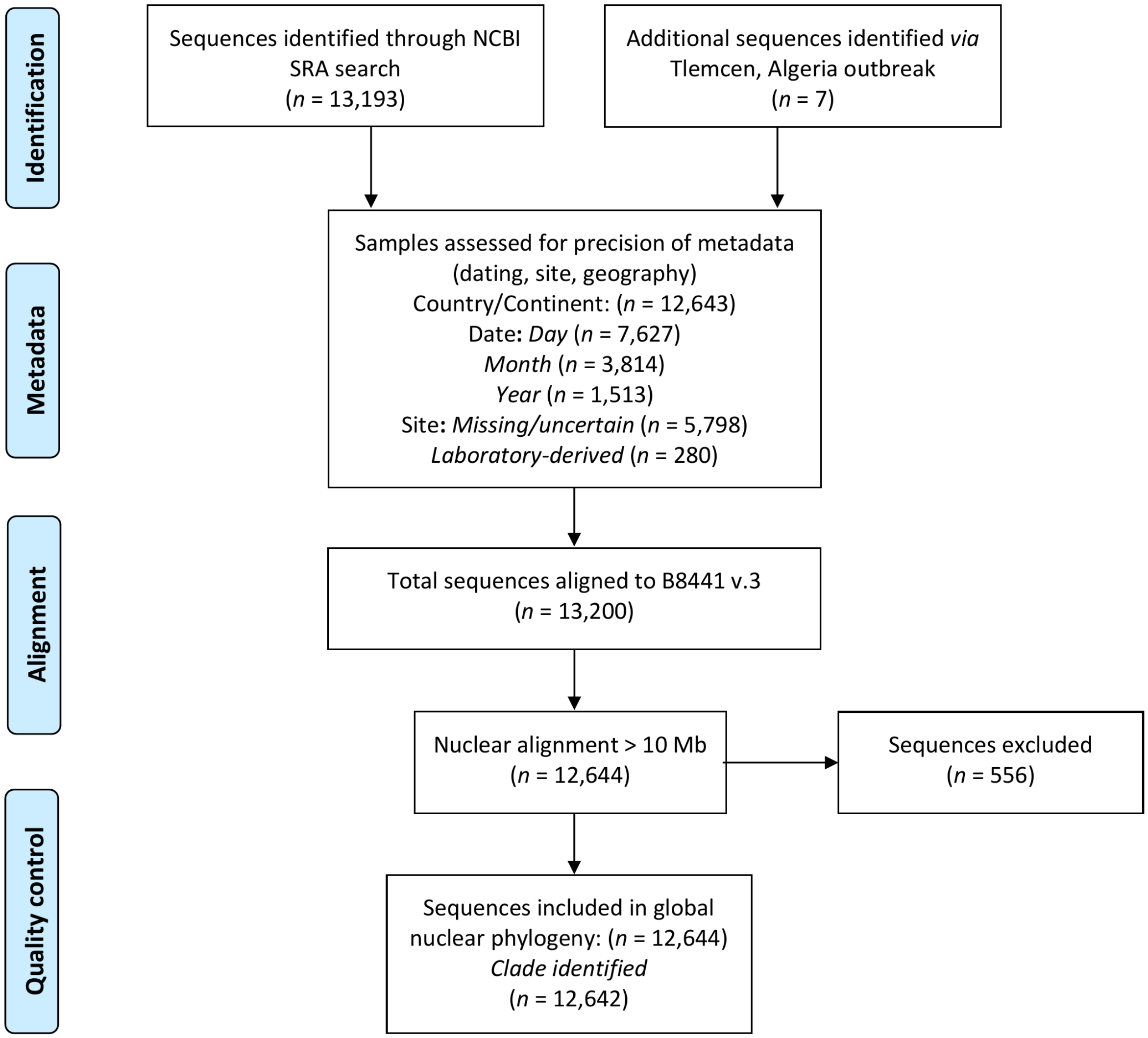


Figure S1: Systematic global genomic data analysis flowsheet. Adapted from the Preferred Reporting Items for Systematic Reviews and Meta-Analyses (PRISMA) flow diagram[4].

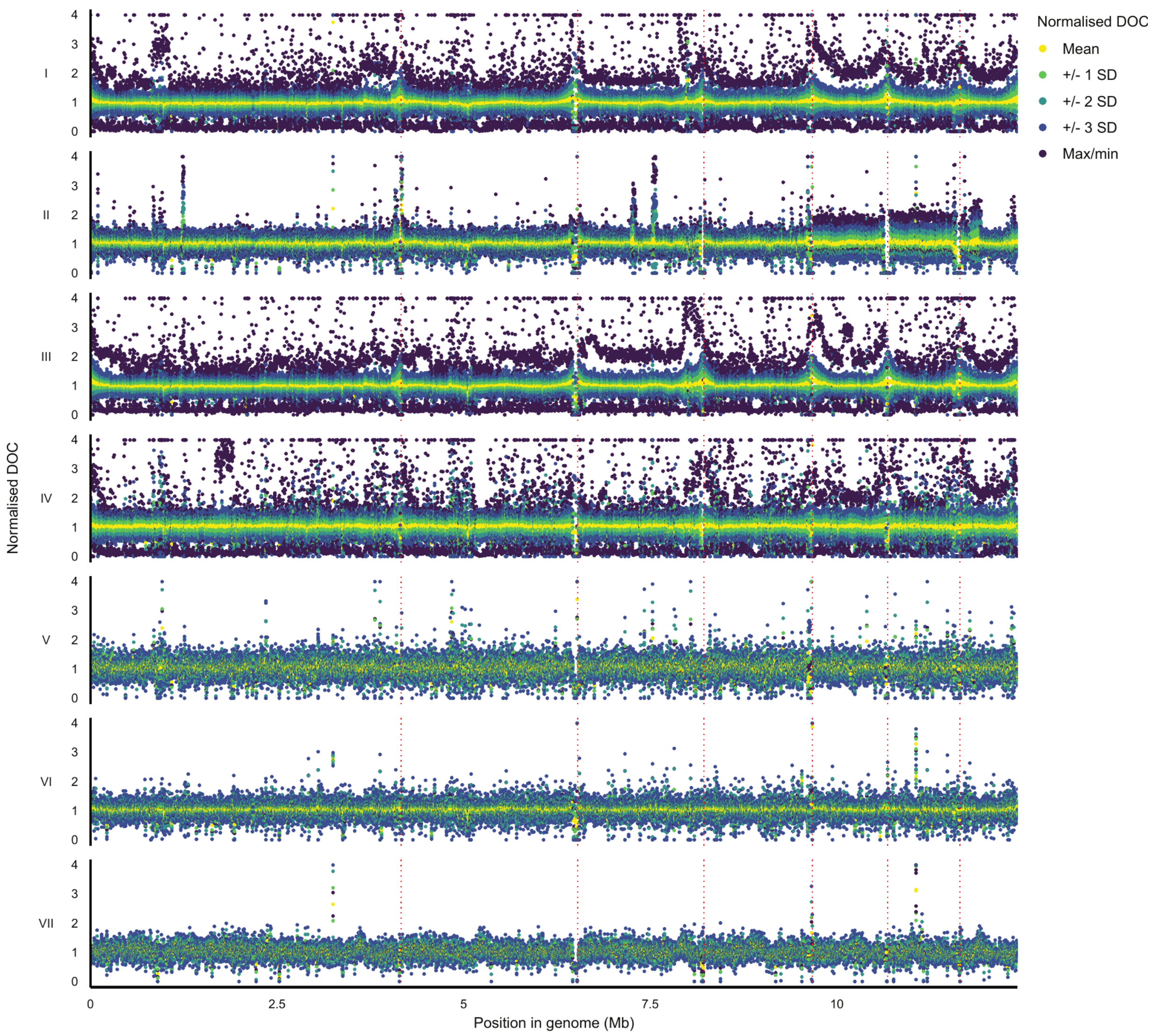


Figure S2: Global copy number variation (CNV): Normalised depth of coverage (NDOC) for each gene locus across the nuclear genome for clinical/environmentally derived accessions with a ceiling of 4. SD: standard deviation. Contig borders are indicated with red dotted vertical lines.

53

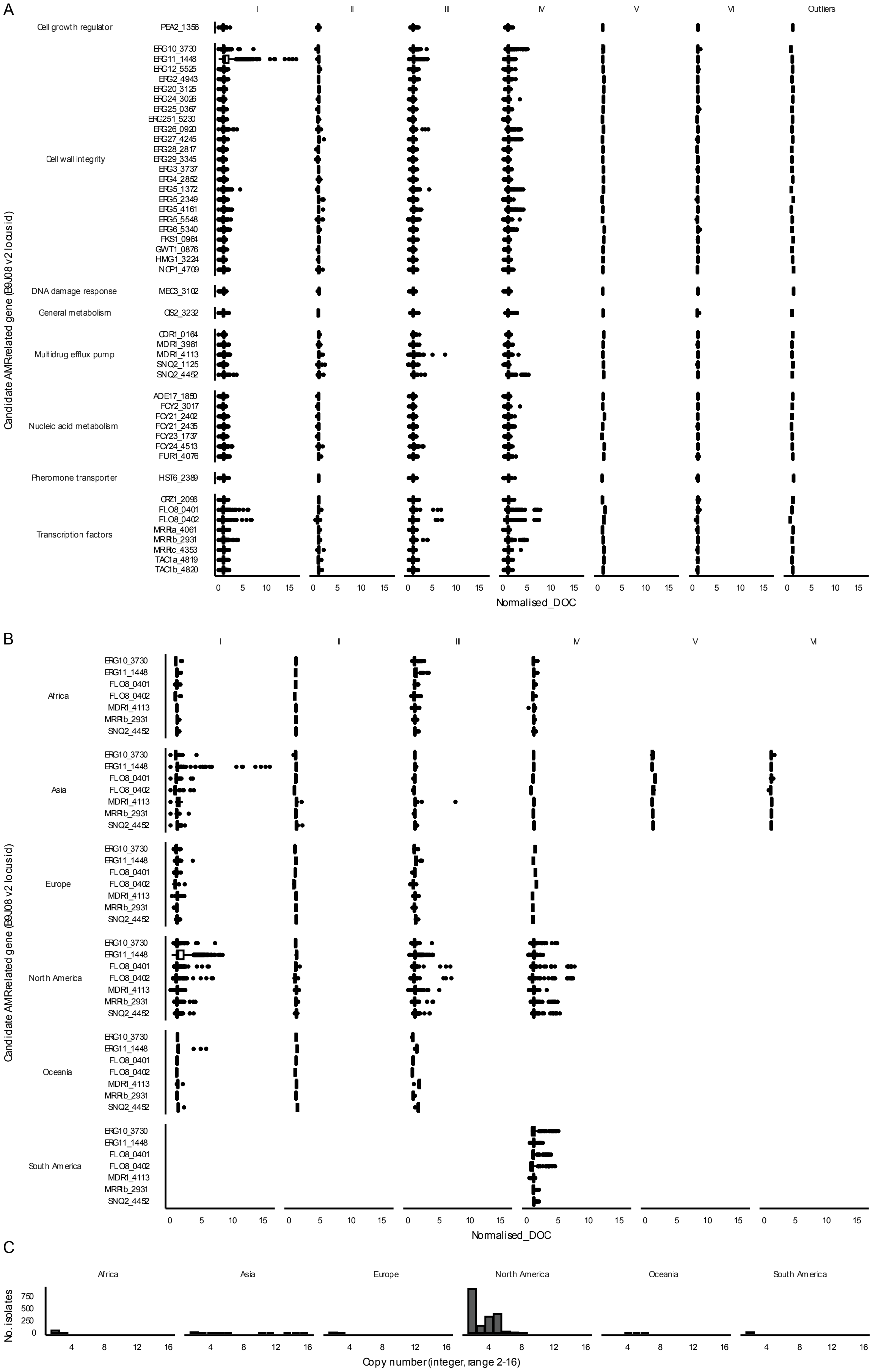

A
I
II
III
IV
V
VI
Outliers
Candidate AMR-related gene (B9J08 v2 locus id)
Cell growth regulator
PEA2_1356
Cell wall integrity
ERG10_3730
ERG11_1448
ERG12_5525
ERG2_4943
ERG20_3125
ERG24_3026
ERG25_0367
ERG251_5230
ERG26_0920
ERG27_4245
ERG28_2817
ERG29_3345
ERG3_3737
ERG4_2852
ERG5_1372
ERG5_2349
ERG5_4161
ERG5_5548
ERG6_5340
FKS1_0964
GWT1_0876
HMG1_3224
NCP1_4709
DNA damage response
MEC3_3102
General metabolism
OS2_3232
Multidrug efflux pump
CDR1_0164
MDR1_3981
MDR1_4113
SNQ2_1125
SNQ2_4452
Nucleic acid metabolism
ADE17_1850
FCY2_3017
FCY21_2402
FCY21_2435
FCY23_1737
FCY24_4513
FUR1_4076
Pheromone transporter
HST6_2389
Transcription factors
CRZ1_2096
FLO8_0401
FLO8_0402
MRR1a_4061
MRR1b_2931
MRR1c_4353
TAC1a_4819
TAC1b_4820
0 5 10 15
Normalised_DOC
B
I
II
III
IV
V
VI
Candidate AMR-related gene (B9J08 v2 locus id)
Africa
Asia
Europe
North America
Oceania
South America
ERG10_3730
ERG11_1448
FLO8_0401
FLO8_0402
MDR1_4113
MRR1b_2931
SNQ2_4452
0 5 10 15
Normalised_DOC
C
Africa
Asia
Europe
North America
Oceania
South America
No. isolates
750
500
250
0
4 8 12 16
Copy number (integer, range 2-16)

Figure S3: Candidate AMR gene copy number variation (CNV): Normalised depth of coverage (NDOC) for clinically/environmentally derived accessions. *(A)*: NDOC across 47 genes for all strains (*n* = 12,369). *(B)*: NDOC across AMR-related genes with NDOC ≥5. *(C)*: Count of *ERG11* genes demonstrating duplication or greater CNV events per continent, illustrating distribution of events.

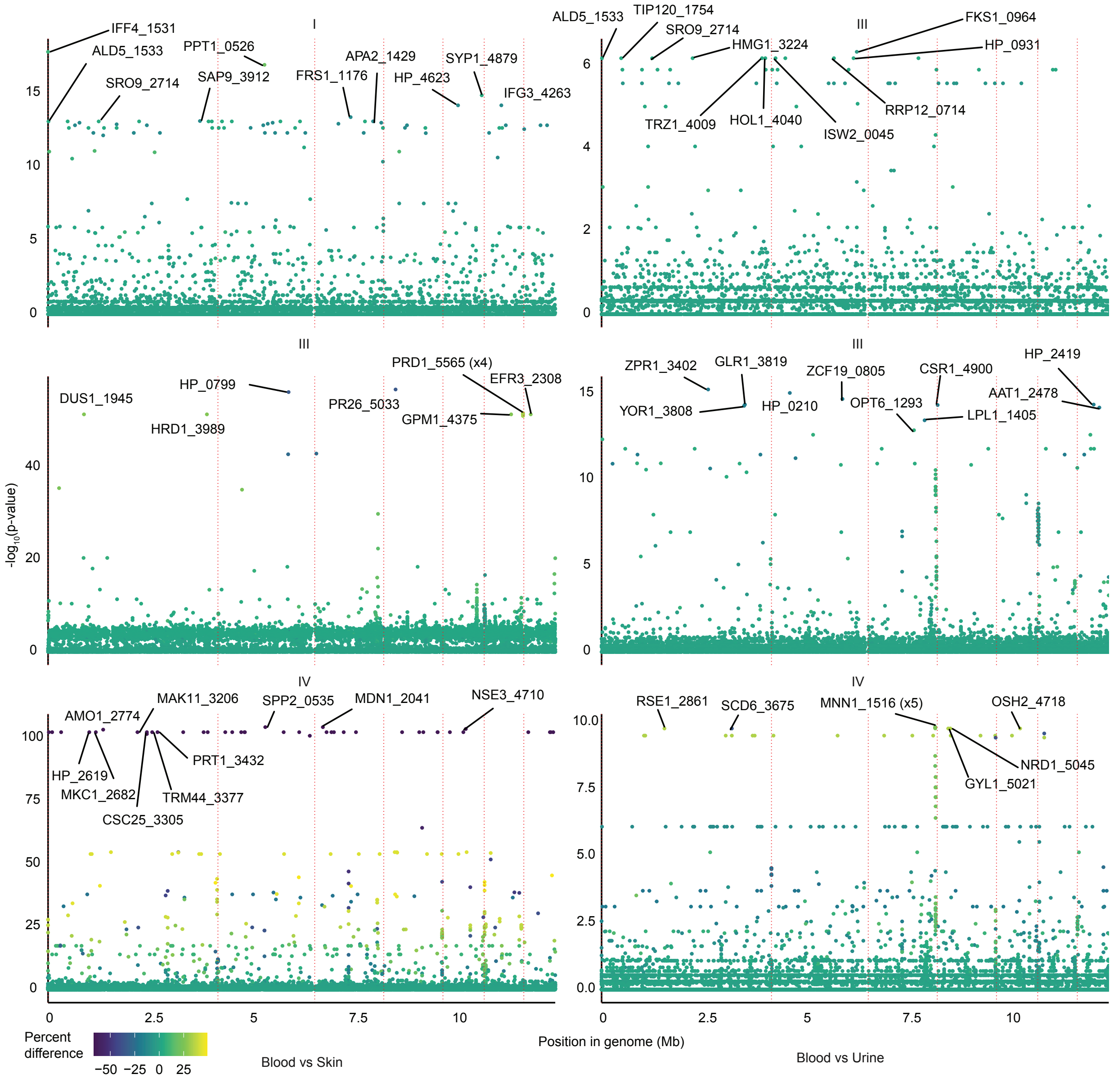

Figure S4: Screening global data for genome-wide association within each clade I, III and IV between epidemiologically derived blood, urine, and skin (axilla/groin) isolates. Ten loci from each comparison with the highest -$\log_{10}$(p-value) after adjusting for multiple correction are labelled.

### Supplementary tables

| BioProject | Number | Regisitered depositing institution | Details | Classification |
|---|---|---|---|---|
| PRJDB6988 | 7 | Laboratory of Bacterial Genomics, Pathogen Genomics Center, National Institute of Infectious Diseases, Japan | Clade II otitis media strains[5] | Epidemiological |
| PRJEB14717 | 3 | Vallabhbhai Patel Chest Institute | Assumed to be linked to early Indian investigation[6] | Epidemiological |
| PRJEB20230 | 27 | Imperial College London, radboudumc | Illumina paired end sequencing of *Candida auris* within the UK | Epidemiological |
| PRJEB21518 | 70 | Wellcome Sanger Institute | First European outbreak in London[7] | Epidemiological |
| PRJEB29190 | 1 | University of Aberdeen | Role of ABC proteins[8] | Laboratory |
| PRJEB34199 | 19 | Public Health Laboratory Centre, Hong Kong | Draft genome sequences Hong Kong[9] | Epidemiological |
| PRJEB36563 | 54 | Imperial College London, radboudumc | King's college outbreak[2] | Epidemiological |
| PRJEB36822 | 65 | Imperial College London, radboudumc | Colonisation isolates from UK outbreak[2] | Epidemiological |
| PRJEB40741 | 20 | University of Aberdeen | *In vitro* evolution of caspofungin resistance in *Candida auris* | Laboratory |
| PRJEB64175 | 1 | National Institute of Health Dr. Ricardo Jorge, Lisbon, Portugal | First case Portugal[10] | Epidemiological |
| PRJEB70513 | 35 | FISABIO, Valencia, Spain | Spanish outbreak[11] | Epidemiological |
| PRJEB9463 | 1 | Vallabhbhai Patel Chest Institute | Early Indian investigation - fluconazole resistant[12] | Epidemiological |
| PRJNA1000034 | 3 | Genome Institute of Singapore | Clade VI in Singapore[13] | Epidemiological |
| PRJNA1003896 | 99 | Centers for Disease Control and Prevention | Genetic epidemiology in Colombia[14] | Epidemiological |
| PRJNA1004889 | 4 | University of California Los Angeles | Fungal identification | Epidemiological |
| PRJNA1015380 | 2 | Fudan University, China | Assumed to be linked to[15] | Epidemiological |
| PRJNA1034667 | 250 | Washington State Department of Health | Eligible isolates include clinical isolates associated with invasive infection, as well as isolates recovered from positive colonization screening swabs. | Epidemiological |
| PRJNA1036037 | 10 | University of Lausanne | Assumed to be linked to Fluconazole microevolution[16] | Laboratory |
| PRJNA1039591 | 48 | National Institute for Communicable Diseases, South Africa | *Candida auris* genomic sequence data. | Epidemiological |
| PRJNA1085724 | 340 | Wisconsin State Laboratory of Hygiene | *Candida auris* WGS | Epidemiological |
| PRJNA1092051 | 420 | Texas Department of State Health Services | Whole Genome Sequencing of *Candida auris* | Epidemiological |
| PRJNA1096176 | 3 | IRB Barcelona | Evolved amphotericin B resistance[17,18] | Laboratory |
| PRJNA1096699 | 8 | University of Exeter | Clinical strains interrogated for adhesion dynamics[19] | Laboratory |
| PRJNA1134646 | 333 | Maryland Department of Health Core Sequencing Laboratory submission group | CDC Mycotic Diseases Branch Candida auris pathogen surveillance | Epidemiological |
| PRJNA267757 | 3 | Indian Institute of Science (IISc) | Early strain assembly[20] | Epidemiological |

| | | | | |
|---|---|---|---|---|
| PRJNA328792 | 51 | National Center for Emerging and Zoonotic Infectious Diseases- Mycotic Diseases Branch | Simultaneous emergence[21], genomic insights into MDR/mating[22], and population genetic tracing[23] | Epidemiological |
| PRJNA415955 | 78 | University of Oxford | John Radcliffe Hospital outbreak[24] | Epidemiological |
| PRJNA470683 | 82 | National Center for Emerging and Zoonotic Infectious Diseases- Mycotic Diseases Branch | Epidemiology in Colombia[25], population genetic tracing[23] | Epidemiological |
| PRJNA480539 | 1 | University of California San Francisco/Chan Zuckerberg Biohub | Returning traveller case (India)[26] | Epidemiological |
| PRJNA485022 | 1 | Murdoch University, Australia | Sternal osteomyelitis case in traveller from Kenya to Australia[27] | Epidemiological |
| PRJNA485145, PRJNA485239, PRJNA485409, PRJNA485414, PRJNA485415 | 5 | Bioinformatik, Biozentrum, Uni Wuerzburg | Cases in Germany linked to overseas healthcare[28] | Epidemiological |
| PRJNA493622 | 315 | National Center for Emerging and Zoonotic Infectious Diseases- Mycotic Diseases Branch | Molecular epidemiology of introductions into the USA[29]; population genetic tracing[23] | Epidemiological |
| PRJNA514082 | 1 | North-Western State Medical University named after I.I. Mechnicov, Russia | First case in Russia[30] | Epidemiological |
| PRJNA540907 | 7 | Singapore General Hospital | Genomic epidemiology in Singapore[31] | Epidemiological |
| PRJNA540998 | 4 | Houston Methodist Hospital, Texas | Patient-colelcted[32] | Epidemiological |
| PRJNA541007 | 1 | National Center for Emerging and Zoonotic Infectious Diseases- Mycotic Diseases Branch | Potential fifth clade[33] | Epidemiological |
| PRJNA549344 | 98 | Institute of Microbiology Chinese Academy of Sciences | Genomic epidemiology in Shenyang, China[34] | Epidemiological |
| PRJNA549561 | 29 | Institute of Microbiology Chinese Academy of Sciences | Adaptive aneuploidy in azole resistance[35] | Laboratory |
| PRJNA560710 | 2 | Canisius-Wilhelmina Hospital, Nijmegen | First two cases in Netherlands[36] | Epidemiological |
| PRJNA592373 | 27 | Public Health Agency of Canada | Screening Canadian patients[37,38] - note 5.7% prevalence after Indian care, and all 4 clades identified | Epidemiological |
| PRJNA595978 | 154 | National Center for Emerging and Zoonotic Infectious Diseases- Mycotic Diseases Branch | Population genetic tracing[23] | Epidemiological |
| PRJNA603602 | 18 | Institute of Clinical Pathology and Medical Research, Westmead Hospital, Australia | Assumed to be linked to PRJNA559200 Australian screening[39] | Epidemiological |
| PRJNA635156 | 17 | University of Minnesota | Clinical strains for *in vitro* evolution of *Candida auris* clinical isolates in 1 μg/ml fluconazole for 30 generations[40] | Laboratory |
| PRJNA635167 | 98 | University of Minnesota | *In vitro* evolution of *Candida auris* clinical isolates in 1 μg/ml fluconazole for 30 generations[40] | Laboratory |
| PRJNA638416 | 1695 | National Center for Emerging and Zoonotic Infectious Diseases- Mycotic Diseases Branch | *Candida auris* genomics surveillance in the United State | Epidemiological |
| PRJNA640677 | 23 | NY Public Health's Wadsworth Center Mycology Laboratory | Pan-resistant strain[41] | Epidemiological |
| PRJNA655187 | 60 | University of Genoa | Genomic epidemiology in Italy[42] and *in vivo* microevolution of echinocandin resistance[43] | Epidemiological |

| | | | | |
|---|---|---|---|---|
| PRJNA664007 | 6 | KU Leuven | Evolved antifungal resistance[17,44] | Laboratory |
| PRJNA672695 | 6 | University of California Los Angeles | Clade III in California[45] | Epidemiological |
| PRJNA672955 | 4 | NHGRI/NIH, Bethesda | *Candida auris* colonization on a ventilator ward[46] | Epidemiological |
| PRJNA679832 | 13 | McMaster University and Department of Medical Mycology, Vallabhbhai Patel Chest Institute, University of Delhi, India | Andaman Islands[47] | Epidemiological |
| PRJNA686013 | 1 | University of Messina, Italy | Egyptian case assumed to be linked to[48] | Epidemiological |
| PRJNA687539 | 9 | McMaster University - Department of Medical Mycology, Vallabhbhai Patel Chest Institute, University of Delhi, Delhi, India | Respiratory *C. auris* in Delhi[49] | Epidemiological |
| PRJNA693430 | 122 | Sidra Medicine, Qatar | Genomic epidemiology of *C. auris* in Qatat[50] including multiple drug resistance[51] | Epidemiological |
| PRJNA722434 | 59 | National Center for Emerging and Zoonotic Infectious Diseases- Mycotic Diseases Branch | Global *Candida auris* genomics surveillance | Epidemiological |
| PRJNA722500 | 2 | University of Michigan | Informing laboratory mutations[52] | Laboratory |
| PRJNA732280 | 1 | University of Michigan | *Candida auris* strain AR0390 from the CDC/FDA antibiotic resistance bank. | Laboratory |
| PRJNA736342 | 1 | UW-Madison | Part of Y1000 evolutionary project[53] | Laboratory |
| PRJNA737309 | 115 | National Institute for Communicable Diseases, South Africa | Clade distribution in South Africa[1] with follow-up *in vitro* studies[54,55] | Epidemiological |
| PRJNA772662 | 6 | UF Health Jacksonville | Not given. Could be linked to[56] | Epidemiological |
| PRJNA792028 | 7 | Fudan University, China | Clinical isolates' amplification of *ALS4*[57] | Epidemiological |
| PRJNA796037 | 80 | National Center for Emerging and Zoonotic Infectious Diseases- Mycotic Diseases Branch | Transmission in post-acute care settings in OC[58] | Epidemiological |
| PRJNA809768 | 16 | McMaster University and University of Delhi | *C. auris* on picked apples[59] | Epidemiological |
| PRJNA816104 | 4 | Canisius-Wilhelmina Hospital, Nijmegen | Clade V isolates[60] | Epidemiological |
| PRJNA838244 | 4 | African Centre of Excellence for Genomics of Infectious Diseases (ACEGID), Redeemer's University | Whole genome sequencing of *C. auris* from patients presenting with fungaemia in Lagos State, Nigeria[61] | Epidemiological |
| PRJNA843706 | 24 | University of Szeged | Triazole microevolution[62] | Laboratory |
| PRJNA846332 | 2752 | Nevada State Public Health Laboratory submission group | Detection of opposite mating types[63]. Assumed to be linked to[64] | Epidemiological |
| PRJNA847553 | 2987 | Utah Public Health Laboratory Infectious Disease submission group | Genome sequencing and assembly of *Candida auris* species at UPHL. | Epidemiological |
| PRJNA848263 | 3 | National University Hospital, Singapore | Singapore isolates[65] | Epidemiological |
| PRJNA856410 | 19 | Microbiological Diagnostic Unit Public Health Laboratory (MDU PHL) Victoria, Australia | Assumed to be linked to Australian outbreak[66] | Epidemiological |
| PRJNA857686 | 236 | Nevada State Public Health Laboratory submission group | GAMBIT (Genomic Approximation Method for Bacterial Identification and Tracking)[67]. Assumed to be linked to[64] | Epidemiological |
| PRJNA863457 | 1 | Nicolaus Copernicus in Torun, Ludwik Rydygier Collegium Medicum in Bydgoszcz, Poland | Case of meningococcal septicaemia[68] | Epidemiological |

| | | | | |
|---|---|---|---|---|
| PRJNA865346 | 2 | Worcester Polytechnic Institute | Correlative study of virulence traits[69] | Laboratory |
| PRJNA865936 | 4 | Institut Pasteur | First in-France nosocomial transmission[70] | Epidemiological |
| PRJNA867138 | 1 | Academy of Military Medical Sciences, China | Case report[71] | Epidemiological |
| PRJNA883226 | 28 | Centers for Disease Control and Prevention | Candida auris samples collected from US states | Epidemiological |
| PRJNA904262 | 2 | University of Michigan | *SCF1* study[72] | Laboratory |
| PRJNA904373 | 618 | Regional Innovative Public Health Laboratory, Chicago, IL | *Candida auris* whole genome sequencing by the Regional Innovative Public Health Laboratory, Chicago, IL. Not given but could be linked to[73] | Epidemiological |
| PRJNA912376 | 16 | Peking Union Medical College Hospital | Isolates from patients that are *in vivo* evolved[74] | Epidemiological |
| PRJNA917187 | 7 | McMaster University | Isolation from dog ears[75] | Epidemiological |
| PRJNA918317 | 515 | Wadsworth Center, NY State Dept. of health | Wadsworth Center, NY State Dept. of Health *C. auris* WGS | Epidemiological |
| PRJNA923734 | 5 | Medical University of Vienna | Austrian outbreak[76] | Epidemiological |
| PRJNA932032 | 24 | University of California San Francisco | Genome sequencing of clioquinol-evolved *C. auris* strains[77] | Laboratory |
| PRJNA950111 | 1 | Nevada State Public Health Laboratory submission group | Fungus sequenced at Nevada State Public Health Lab that are not *Candida auris*. Assumed to be linked to[64] | Epidemiological |
| PRJNA979221 | 430 | Orange County Public Health Lab | *Candida auris* sequences by the Orange County Public Health Lab | Epidemiological |
| PRJNA981511 | 10 | National Center for Emerging and Zoonotic Infectious Diseases (NCEZID) and ICDDR Bangladesh | Clade VI emergence Bangladesh[78] | Epidemiological |
| PRJNA982799 | 32 | Canisius-Wilhelmina Hospital, Nijmegen | Kuwait outbreak[79] | Epidemiological |
| PRJNA984918 | 32 | KU Leuven | Evolved amphotericin B resistance[17,18] | Laboratory |
| PRJNA988017 | 253 | Minnesota Department of Health Infectious Disease Laboratory Submission Group | Candida auris datasets from clinical and environmental samples applicable for molecular epidemiology. | Epidemiological |
| PRJNA991119, PRJNA1035021 | 23 | National Institute for Infectious Diseases Matei Bals, Romania | Romanian outbreak[80] | Epidemiological |
| PRJNA991735 | 118 | National Health Laboratory Service, South Africa | Likely to be part of[81] | Epidemiological |
| PRJNA999713 | 1 | National Center for Emerging and Zoonotic Infectious Diseases- Mycotic Diseases Branch | Dog ear Kansas[82] | Epidemiological |

Table S1: Global genomic epidemiology sequencing projects.

| Accession | Sample type | Collection date | Clade (this study) | Clade (STR) | *ERG11* | *FKS1* |
|---|---|---|---|---|---|---|
| L0048 | Urine | 2017-09-01 | II | II | - | L1572I |
| L0049 | Bronchioalveolar lavage | 2017-05-26 | IV | IV | Y132F, K177R, N335S, E343D | - |
| L0050 | Ascites | 2018-08-04 | II | IV | - | L1572I |
| L0051 | Wound | 2019-07-26 | IV | IV | Y132F, K177R, N335S, E343D | - |
| L0052 | Bronchioalveolar lavage | 2019-08-08 | III | III | - | - |
| L0053 | Bronchioalveolar lavage | 2019-08-02 | III | III | - | - |
| L0054 | Bronchioalveolar lavage | 2018-01-14 | I | I | Y132F | - |

Table S2: Summary features of Algerian isolate whole genome sequences.

### Supplementary references


1. Naicker, S. D. *et al.* Clade distribution of *Candida auris* in South Africa using whole genome sequencing of clinical and environmental isolates. *Emerging Microbes & Infections* **10**, 1300–1308 (2021).
2. Kappel, D. *et al.* Genomic epidemiology describes introduction and outbreaks of antifungal drug-resistant *Candida auris*. *npj Antimicrobials and Resistance* **2**, 1–10 (2024).
3. Zerrouki, H. *et al.* Emergence of *Candida auris* in intensive care units in Algeria. *Mycoses* e13470 (2022) doi:10.1111/myc.13470.
4. Moher, D., Liberati, A., Tetzlaff, J., Altman, D. G. & Group, T. P. Preferred Reporting Items for Systematic Reviews and Meta-Analyses: The PRISMA Statement. *PLOS Medicine* **6**, e1000097 (2009).
5. Sekizuka, T. *et al.* Clade II *Candida auris* possess genomic structural variations related to an ancestral strain. *PLOS ONE* **14**, e0223433 (2019).
6. Sharma, C., Kumar, N., Pandey, R., Meis, J. F. & Chowdhary, A. Whole genome sequencing of emerging multidrug resistant *Candida auris* isolates in India demonstrates low genetic variation. *New Microbes and New Infections* **13**, 77–82 (2016).
7. Schelenz, S. *et al.* First hospital outbreak of the globally emerging *Candida auris* in a European hospital. *Antimicrobial Resistance & Infection Control* **5**, 35 (2016).
8. Wasi, M. *et al.* ABC transporter genes show upregulated expression in drug-resistant clinical isolates of *Candida auris*: A genome-wide characterization of ATP-binding cassette (ABC) transporter genes. *Frontiers in Microbiology* **10**, 1445 (2019).

9. Tse, H., Tsang, A. K. L., Chu, Y.-W. & Tsang, D. N. C. Draft Genome Sequences of 19 Clinical Isolates of *Candida auris* from Hong Kong. *Microbiology Resource Announcements* **10**, e00308–20 (2021).

10. Henriques, J. *et al. Candida auris* in intensive care setting: The first case reported in Portugal. *Journal of Fungi* **9**, 837 (2023).

11. Mulet-Bayona, J. V. *et al.* Genotypic and phenotypic characterisation of a nosocomial outbreak of *Candida auris* in Spain during 5 years. *Mycoses* **67**, e13776 (2024).

12. Sharma, C., Kumar, N., Meis, J. F., Pandey, R. & Chowdhary, A. Draft genome sequence of a fluconazole-resistant *Candida auris* strain from a candidemia patient in India. *Genome Announcements* **3**, e00722–15 (2015).

13. Suphavilai, C. *et al.* Detection and characterisation of a sixth *Candida auris* clade in Singapore: A genomic and phenotypic study. *Lancet Microbe* **5**, (2024).

14. Misas, E. *et al.* Genomic epidemiology and antifungal-resistant characterization of *Candida auris*, Colombia, 2016–2021. *mSphere* **9**, e00577–23 (2024).

15. Bing, J. *et al. Candida auris*-associated hospitalizations and outbreaks, China, 2018–2023. *Emerging Microbes & Infections* **13**, 2302843 (2024).

16. Louvet, M. *et al.* Ume6-dependent pathways of morphogenesis and biofilm formation in *Candida auris*. *Microbiology Spectrum* **2**, e01531–24 (2024).

17. Carolus, H. *et al.* Acquired amphotericin B resistance leads to fitness trade-offs that can be mitigated by compensatory evolution in *Candida auris*. *Nature Microbiology* **9**, 3304–3320 (2024).

18. Carolus, H. *et al.* Collateral sensitivity counteracts the evolution of antifungal drug resistance in *Candida auris*. *Nature Microbiology* **9**, 2954–2969 (2024).

19. Malavia-Jones, D. *et al.* Strain and temperature dependent aggregation of *Candida auris* is attenuated by inhibition of surface amyloid proteins. *The Cell Surface* **10**, 100110 (2023).

20. Chatterjee, S. *et al.* Draft genome of a commonly misdiagnosed multidrug resistant pathogen *Candida auris*. *BMC Genomics* **16**, 686 (2015).

21. Lockhart, S. R. *et al.* Simultaneous emergence of multidrug-resistant *Candida auris* on 3 continents confirmed by whole-genome sequencing and epidemiological analyses. *Clinical Infectious Diseases* **64**, 134–140 (2017).

22. Muñoz, J. F. *et al.* Genomic insights into multidrug-resistance, mating and virulence in *Candida auris* and related emerging species. *Nature Communications* **9**, 5346 (2018).

23. Chow, N. A. *et al.* Tracing the evolutionary history and global expansion of *Candida auris* using population genomic analyses. *mBio* **11**, 15 (2020).

24. Eyre, D. W. *et al.* A *Candida auris* outbreak and its control in an intensive care setting. *New England Journal of Medicine* **379**, 1322–1331 (2018).

25. Escandón, P. *et al.* Molecular epidemiology of *Candida auris* in Colombia reveals a highly related, countrywide colonization with regional patterns in amphotericin B resistance. *Clinical Infectious Diseases* **68**, 15–21 (2019).
26. Woodworth, M. H. *et al.* Sentinel case of *Candida auris* in the western United States following prolonged occult colonization in a returned traveler from India. *Microbial Drug Resistance* **25**, 677–680 (2019).
27. Heath, C. H., Dyer, J. R., Pang, S., Coombs, G. W. & Gardam, D. J. *Candida auris* sternal osteomyelitis in a man from Kenya visiting Australia, 2015. *Emerging Infectious Diseases* **25**, 192–194 (2019).
28. Hamprecht, A. *et al. Candida auris* in Germany and previous exposure to foreign healthcare. *Emerging Infectious Diseases* **25**, 1763–1765 (2019).
29. Chow, N. A. *et al.* Multiple introductions and subsequent transmission of multidrug-resistant *Candida auris* in the USA: A molecular epidemiological survey. *The Lancet. Infectious diseases* **18**, 1377–1384 (2018).
30. Pchelin, I. M. *et al.* Whole genome sequence of first *Candida auris* strain, isolated in Russia. *Medical Mycology* **58**, 414–416 (2020).
31. Tan, Y. E. *et al. Candida auris* in Singapore: Genomic epidemiology, antifungal drug resistance, and identification using the updated 8.01 VITEK(r)2 system. *International Journal of Antimicrobial Agents* **54**, 709–715 (2019).
32. Long, S. W., Olsen, R. J., Nguyen, H. A. T., Saavedra, M. O. & Musser, J. M. Draft genome sequence of *Candida auris* strain LOM, a human clinical isolate from greater metropolitan Houston, Texas. *Microbiology Resource Announcements* **8**, 3 (2019).
33. Chow, N. A. *et al.* Potential fifth clade of *Candida auris,* Iran, 2018. *Emerging Infectious Diseases* **25**, 1780–1781 (2019).
34. Tian, S. *et al.* Genomic epidemiology of *Candida auris* in a general hospital in Shenyang, China: A three-year surveillance study. *Emerging Microbes & Infections* **10**, 1088–1096 (2021).
35. Bing, J. *et al.* Experimental evolution identifies adaptive aneuploidy as a mechanism of fluconazole resistance in *Candida auris*. *Antimicrobial Agents and Chemotherapy* **65**, e01466–20 (2020).
36. Vogelzang, E. H. *et al.* The first two cases of *Candida auris* in The Netherlands. *Journal of Fungi* **5**, 91 (2019).
37. Garcia-Jeldes, H. F. *et al.* Prevalence of *Candida auris* in Canadian acute care hospitals among at-risk patients, 2018. *Antimicrobial Resistance & Infection Control* **9**, 82 (2020).
38. De Luca, D. G. *et al.* Four genomic clades of *Candida auris* identified in Canada, 2012–2019. *Medical Mycology* **60**, myab079 (2022).

39. Biswas, C. *et al.* Genetic heterogeneity of Australian *Candida auris* isolates: Insights from a nonoutbreak setting using whole-genome sequencing. *Open Forum Infectious Diseases* **7**, ofaa158 (2020).

40. Burrack, L. S., Todd, R. T., Soisangwan, N., Wiederhold, N. P. & Selmecki, A. Genomic diversity across *Candida auris* clinical isolates shapes rapid development of antifungal resistance *in vitro* and *in vivo*. *mBio* **13**, e0084222 (2022).

41. Jacobs, S. E. *et al. Candida auris* pan-drug-resistant to four classes of antifungal agents. *Antimicrobial Agents and Chemotherapy* **66**, e0005322 (2022).

42. Di Pilato, V. *et al.* Molecular epidemiological investigation of a nosocomial cluster of *C. auris*: Evidence of recent emergence in Italy and ease of transmission during the COVID-19 pandemic. *Journal of Fungi* **7**, 140 (2021).

43. Codda, G. *et al. In vivo* evolution to echinocandin resistance and increasing clonal heterogeneity in *Candida auris* during a difficult-to-control hospital outbreak, Italy, 2019 to 2022. *Eurosurveillance* **28**, 2300161 (2023).

44. Carolus, H. *et al.* Genome-wide analysis of experimentally evolved *Candida auris* reveals multiple novel mechanisms of multidrug resistance. *mBio* **12**, e03333–20 (2021).

45. Price, T. K. *et al.* Genomic characterizations of clade III lineage of *Candida auris*, California, USA. *Emerging Infectious Diseases* **27**, 1223–1227 (2021).

46. Proctor, D. M. *et al.* Integrated genomic, epidemiologic investigation of *Candida auris* skin colonization in a skilled nursing facility. *Nature Medicine* **27**, 1401–1409 (2021).

47. Arora, P. *et al.* Environmental isolation of *Candida auris* from the coastal wetlands of Andaman Islands, India. *mBio* **12**, e03181–20 (2021).

48. El-Kholy, M., Shawky, S., Fayed, A. & Meis, J. *Candida auris* bloodstream infection in Egypt. In: 9th Trends in Medical Mycology held on 11–14 October 2019, Nice, France, organized under the auspices of EORTC-IDG and ECMM. *Journal of Fungi* **5**, 95 (2019).

49. Yadav, A. *et al.* Colonisation and transmission dynamics of *Candida auris* among chronic respiratory diseases patients hospitalised in a chest hospital, Delhi, India: A comparative analysis of whole genome sequencing and microsatellite typing. *Journal of Fungi* **7**, 81 (2021).

50. Salah, H. *et al.* Genomic epidemiology of *Candida auris* in Qatar reveals hospital transmission dynamics and a South Asian origin. *Journal of Fungi* **7**, 240 (2021).

51. Abid, F. B. *et al.* Molecular characterization of *Candida auris* outbreak isolates in Qatar from patients with COVID-19 reveals the emergence of isolates resistant to three classes of antifungal drugs. *Clinical Microbiology and Infection* **29**, 1083.e1–1083.e7 (2023).

52. Santana, D. J. & O'Meara, T. R. Forward and reverse genetic dissection of morphogenesis identifies filament-competent *Candida auris* strains. *Nature Communications* **12**, 7197 (2021).

53. Shen, X.-X. *et al.* Tempo and mode of genome evolution in the budding yeast subphylum. *Cell* **175**, 1533–1545.e20 (2018).

54. Maphanga, T. G. *et al. In vitro* antifungal resistance of *Candida auris* isolates from bloodstream infections, South Africa. *Antimicrobial Agents and Chemotherapy* **65**, (2021).

55. Maphanga, T. G., Mpembe, R. S., Naicker, S. D., Govender, N. P. & for GERMS-SA. *In vitro* antifungal activity of manogepix and other antifungal agents against South African *Candida auris* isolates from bloodstream infections. *Microbiology Spectrum* **10**, e01717–21 (2022).

56. Goulart, M. A. *et al.* Identification and infection control response to Candida auris at an academic level I trauma center. *American Journal of Infection Control* **52**, 371–373 (2024).

57. Bing, J. *et al.* Clinical isolates of *Candida auris* with enhanced adherence and biofilm formation due to genomic amplification of *ALS4*. *PLOS Pathogens* **19**, e1011239 (2023).

58. Karmarkar, E. N. *et al.* Rapid assessment and containment of *Candida auris* transmission in postacute care settings-Orange County, California, 2019. *Annals of Internal Medicine* **174**, 1554–1562 (2021).

59. Yadav, A. *et al. Candida auris* on apples: Diversity and clinical significance. *mBio* **13**, e00518–22 (2022).

60. Spruijtenburg, B. *et al.* Confirmation of fifth *Candida auris* clade by whole genome sequencing. *Emerging Microbes & Infections* 1–15 (2022) doi:10.1080/22221751.2022.2125349.

61. Oladele, R. *et al.* Emergence and genomic characterization of multidrug resistant *Candida auris* in Nigeria, West Africa. *Journal of Fungi* **8**, 787 (2022).

62. Bohner, F. *et al.* Acquired triazole resistance alters pathogenicity-associated features in *Candida auris* in an isolate-dependent manner. *Journal of Fungi* **9**, 1148 (2023).

63. Massic, L. *et al.* Detection of five instances of dual-clade infections of *Candida auris* with opposite mating types in southern Nevada, USA. *The Lancet Infectious Diseases* **23**, e328–e329 (2023).

64. Gorzalski, A. *et al.* The use of whole-genome sequencing and development of bioinformatics to monitor overlapping outbreaks of *Candida auris in southern Nevada*. *Frontiers in Public Health* **11**, 1198189 (2023).

65. Chew, K. L., Achik, R., Osman, N. H., Octavia, S. & Teo, J. W. P. Genomic epidemiology of human candidaemia isolates in a tertiary hospital. *Microbial Genomics* **9**, mgen001047 (2023).

66. Lane, C. R. *et al.* Incursions of *Candida auris* into Australia, 2018. *Emerging Infectious Diseases* **26**, 1326–1328 (2020).

67. Lumpe, J. *et al.* GAMBIT (Genomic Approximation Method for Bacterial Identification and Tracking): A methodology to rapidly leverage whole genome sequencing of bacterial isolates for clinical identification. *PLOS ONE* **18**, e0277575 (2023).

68. Prażyńska, M. *et al. Candida auris* infection in a meningococcal septicemia survivor, Poland. *Mycopathologia* **188**, 135–141 (2023).

69. Yang, B. *et al.* A correlative study of the genomic underpinning of virulence traits and drug tolerance of *Candida auris*. *Infection and Immunity* **92**, e00103–24 (2024).

70. Alanio, A. *et al.* First patient-to-patient intrahospital transmission of clade I *Candida auris* in France revealed after a two-month incubation period. *Microbiology Spectrum* **10**, e01833–22 (2022).

71. Xu, Z. *et al.* A candidemia case caused by a novel drug-resistant *Candida auris* with the Y132F mutation in Erg11 in mainland China. *Infection and Drug Resistance* **16**, 3065–3072 (2023).

72. Santana, D. J. *et al.* A *Candida auris*-specific adhesin, Scf1, governs surface association, colonization, and virulence. *Science* **381**, 1461–1467 (2023).

73. Mitchell, B. I. *et al.* Identifying *Candida auris* transmission in a hospital outbreak investigation using whole-genome sequencing and SNP phylogenetic analysis. *Journal of Clinical Microbiology* **2**, e00680–24 (2024).

74. Chen, X.-F. *et al.* Genome-wide analysis of in vivo-evolved *Candida auris* reveals multidrug-resistance mechanisms. *Mycopathologia* **189**, 35 (2024).

75. Yadav, A. *et al.* *Candida auris* in dog ears. *Journal of Fungi* **9**, 720 (2023).

76. Spettel, K. *et al.* *Candida auris* in Austria—what is new and what is different. *Journal of Fungi* **9**, 129 (2023).

77. Lohse, M. B. *et al.* Broad susceptibility of *Candida auris* strains to 8-hydroxyquinolines and mechanisms of resistance. *mBio* **14**, e01376–23 (2023).

78. Khan, T. *et al.* Emergence of the novel sixth *Candida auris* clade VI in Bangladesh. *Microbiology Spectrum* **12**, e03540–23 (2024).

79. Spruijtenburg, B. *et al.* Whole genome sequencing analysis demonstrates therapy-induced echinocandin resistance in *Candida auris* isolates. *Mycoses* **66**, 1079–1086 (2023).

80. Stanciu, A. M. *et al.* First report of *Candida auris* in Romania: Clinical and molecular aspects. *Antimicrobial Resistance and Infection Control* **12**, 91 (2023).

81. Shuping, L. *et al.* High prevalence of *Candida auris* colonization during protracted neonatal unit outbreak, South Africa. *Emerging Infectious Diseases* **29**, 1913–1916 (2023).

82. White, T. C. *et al.* *Candida auris* detected in the oral cavity of a dog in Kansas. *mBio* **15**, e03080–23 (2024).
